# Constraints for spatially and temporally precise learning in a neural circuit model of reinforcement learning

**DOI:** 10.64898/2026.07.22.740062

**Authors:** Yiheng Wang, Joergen Kornfeld, Michale Fee, Mark Goldman

## Abstract

Reinforcement learning is a key means by which animals learn appropriate actions in a given context. A large body of work suggests that such learning depends on interactions between cortico-basal ganglia circuits and the midbrain dopaminergic system, yet the underlying circuit mechanisms and plasticity rules are not fully understood. Here we present a biologically plausible, multi-region neural circuit model of songbird vocal learning and map it onto the actor-critic framework of reinforcement learning. In this model, stochastic spiking activity in the cortico-basal ganglia pathway implements action selection and drives behavioral exploration, while the pathways driving midbrain dopaminergic signaling evaluate behavioral outcomes and support a reward prediction error based learning rule that approximates stochastic gradient ascent. The model achieves millisecond-scale precise learning that matches observed behavior. We further use the model to examine two fundamental constraints on biological reinforcement learning. First, dopaminergic reinforcement signals are temporally imprecise, which can cause interference between neurons controlling actions that occur close in time. Second, dopaminergic signals are spatially imprecise, which can cause interference between neurons controlling different aspects of behavior but receiving a common reinforcement signal. By jointly modeling the actor and critic components of the circuit, we show that fast updating of reward prediction is crucial for precise and efficient learning under both forms of interference, and the model predicts the experimentally observed timescale of reward prediction updating. These results suggest a circuit-level mechanism by which biological systems achieve reinforcement learning despite the temporal and spatial limitations of global neuromodulatory signals.

## 1 Introduction

Reward-based learning is a cornerstone of adaptive behaviors across species, from animals selecting actions that lead to rewards [1, 2] to humans mastering complex skills like cooking and swimming. Trial-and-error learning [3] has long been recognized as a key mechanism for solving such problems, with the underlying algorithms provided by the theoretical field of reinforcement learning [4]. Neural substrates of reinforcement learning have been widely identified in the brain [5, 6]. For example, cortico-basal ganglia circuits have been implicated in action selection [7, 8, 9, 10, 11] and biological reinforcement learning [12, 13, 14], whereas dysfunction of these circuits has been implicated in psychiatric disorders [15, 16]. The substantia nigra pars compacta (SNc) and the ventral tegmental area (VTA) have been shown in many species to produce dopamine signals that resemble the reward prediction errors (RPEs) that are crucial for reinforcement learning [17, 18, 19, 20]. However, the precise neural circuits and plasticity rules that implement reinforcement learning algorithms in the brain remain elusive.

Songbird vocal learning is an ideal animal model for studying biological reinforcement learning [21, 22]. First, song learning has a clear behavioral paradigm [23]. A male songbird learns to sing a mating song to attract female companions. He first listens to a tutor song from his father and memorizes the tutor song as an internal template. He then learns to mimic the tutor song by comparing his own practice song with the template and making adjustments accordingly. Second, the songbird brain has simple circuitry (Figure 1A) highly specialized for song learning [24, 25, 26]. Earlier conceptual models [27] and recent experimental results [28] suggest that the cortico-basal ganglia circuits in songbirds together with the VTA dopaminergic system perform trial-and-error song learning, analogous to the actor-critic framework of reinforcement learning [29, 30]. In this framework, an actor produces exploratory actions according to a behavioral policy and learns to optimize this policy to maximize the reward it receives. A critic learns to predict the expected reward under the current policy and compares the actual reward with its prediction, generating a reward prediction error that guides the critic to update its prediction and the actor to update its policy.

**Figure 1:**
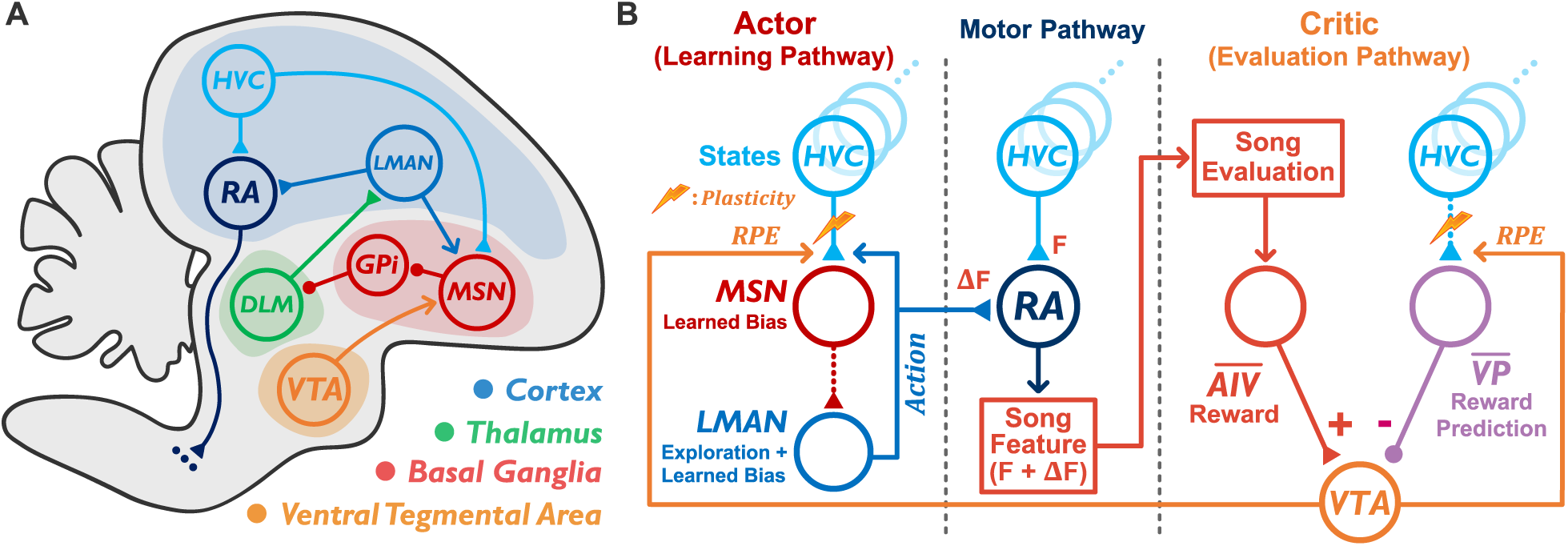
Brain circuitry of song learning. **(A)** A schematic illustration of the brain areas involved in song learning and their known anatomical connections. Triangle and circle heads represent excitatory and inhibitory connections, respectively. Arrowheads (VTA to MSN and LMAN to MSN) represent signals guiding synaptic plasticity. **(B)** The actor-critic neural circuit model. The MSN-LMAN and 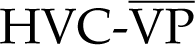 connections are simplified as an effective monosynaptic excitatory connection, indicated by dashed lines. HVC: used as a proper name; RA: robust nucleus of the arcopallium; LMAN: lateral magnocellular nucleus of the anterior nidopallium; MSN: medium spiny neuron; GPi: pallidal-like structure analogous to the mammalian internal globus pallidus; DLM: medial portion of the dorsolateral thalamus; VTA: ventral tegmental area; AIV: ventral intermediate arcopallium; VP: ventral pallidum.

Song production in songbirds involves two distinct pathways: a song motor pathway and a learning pathway. In the motor pathway (Figure 1B, middle), neurons in cortical area HVC (used as a proper name) exhibit robust sequential bursting activity [31, 32, 33] that activates neurons in the downstream area RA (robust nucleus of the arcopallium) to evoke singing behavior. The learning pathway (Figure 1B, left) comprises a cortico-basal ganglia loop thought to be a substrate of trial-and-error reinforcement learning. In this pathway, neurons in nucleus LMAN (lateral magnocellular nucleus of the anterior nidopallium) fire sparsely and stochastically during singing [34, 35, 36] and project to the RA neurons in the motor pathway, producing variations of RA neuron activities and song features like pitch [37, 38, 39, 40]. During each day of singing, the learning pathway implements modifications that correct vocal errors, analogous to the actor in the actor-critic framework. On a slower timescale, these modifications gradually transfer to the motor pathway [38, 41], thus becoming independent of the learning pathway. In this study, we focus on the reinforcement learning process in the learning pathway.

Song learning is thought to be implemented by a three-factor learning rule [42, 22, 43, 44, 45, 46] in which the convergence of temporal context, behavioral exploration, and song performance evaluation guides synaptic plasticity in the learning pathway [27, 28]. Temporal context signals are conveyed by the HVC neurons. An HVC neuron typically generates a single burst (about 10 ms duration) during every song rendition [31, 32, 33]. Different HVC neurons fire at different time points in the song, forming a complete sequence in time. Each HVC neuron can be considered as controlling one 10 ms unit of the song, which we refer to as a song ‘note’. We also refer to a ‘note’ as a ‘state’, using reinforcement learning terminology. HVC neurons project to the medium spiny neurons (MSNs) in the song-related basal ganglia (Area X). Behavioral exploration signals, generated by LMAN neurons, are conveyed to MSNs via axon collaterals from LMAN neurons [47, 48, 27]. Finally, song performance evaluation is conveyed to MSNs by dopaminergic neurons in VTA in the evaluation pathway that encode a reward prediction error (Figure 1B, right). The convergence of these factors can induce plasticity at the HVC-MSN synapses [49, 50] and lead to improvement of song performance [51, 52, 53]. Learned modifications in the firing of MSNs influence song output through a basal ganglia-thalamo-cortical pathway that projects back to LMAN [24, 25, 26].

Analogous to the critic in the actor-critic framework, the evaluation pathway evaluates the song performance and produces the dopamine signals that convey the reward prediction error [54, 55, 56, 53, 57]. Recent experimental evidence suggests a simple neural circuit motif that can compute the reward prediction error (Figure 1B, right). On the one hand, song performance is evaluated in the auditory cortex and leads to a vocal error signal in AIV (ventral intermediate arcopallium) [58, 54]. AIV neurons then inhibit the dopaminergic neurons in VTA by exciting the VTA GABAergic interneurons [30]. Since error can be considered to be a negative reward, as a functional equivalent of this AIV circuitry, we use AIV to represent positive reward signals that functionally project to the VTA dopaminergic neurons with a positive sign. On the other hand, VP (ventral pallidum) neurons express reward prediction-like activity [59]. We use VP to represent reward prediction signals that functionally project to the VTA dopaminergic neurons with a negative sign. Notably, VP also receives input from VTA [59], suggesting VP as a site at which reward-prediction signals may be updated by VTA RPEs. To produce a temporally specific reward prediction signal, the VP neurons need to receive the temporal context. Although anatomically HVC does not project directly to VP, the temporal context can be conveyed indirectly from the song-related basal ganglia [60], or from the vocal thalamic nucleus Uva (Uvaeformis) that forms a motor loop with HVC and contains temporal information about the song [59, 29]. For simplicity, we represent the temporal context with the term ‘HVC’ for both the learning and evaluation pathways (Figure 1B, right). Finally, the VTA neurons receive inputs from both AIV and VP, enabling them to combine the reward and the reward prediction to generate reward prediction error [30, 59, 29].

Here we present a mechanistic actor-critic neural circuit model that performs song learning. We first propose that the core computation of song learning can be viewed as a sequence of approximately independent binary decision problems. The stochastic spiking or not-spiking of the LMAN neurons in the model represents such binary decisions and drives behavioral exploration. Dopamine-guided synaptic plasticity implements an RPE-based learning rule that approximates stochastic gradient ascent on song performance. This mechanism supports accurate song learning that matches experimental results. We then consider two key problems faced broadly by biological reinforcement learning systems. First, dopamine signaling is temporally imprecise, with a timescale of several tens to a hundred milliseconds [55, 56], much slower than the temporally precise singing behavior that operates at a 5-10 millisecond scale [61]. Second, dopamine signaling is spatially imprecise, as a single reward prediction error influences synaptic plasticity across multiple motor channels. Such imprecisions can cause temporal interference between neurons controlling close-in-time actions and spatial interference between neurons controlling different aspects of behavior. Our model provides a first-principles explanation for the fast speed of updating recently observed in songbird learning and more broadly suggests that fast updating of reward prediction is crucial for precise and efficient reinforcement learning.

## 2 Results

### 2.1 Neural circuit model of song learning

We built an actor-critic neural circuit model of the song learning system (Figure 1B, Methods). In our model, the actor component (HVC, MSNs, and LMAN) represents the learning pathway (Figure 1B, left) and the critic component (song evaluation, HVC, 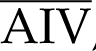, 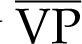, and VTA) represents the evaluation pathway (Figure 1B, right). The motor pathway (Figure 1B, middle) is not explicitly modeled since its activity is stereotyped across the several-hour timescale of learning considered in our model [38, 41]. Instead, we consider the activity along this pathway to represent a fixed contribution to the song feature (*F*), and we model the learned variations (Δ*F*) generated by the actor component around this fixed contribution. In the actor component, stochastic spiking activity of the LMAN neurons, modeled as integrate-and-fire neurons, produce exploratory actions. The firing probabilities of the LMAN neurons at each note of the song can be considered as the combination of a noise-driven baseline exploration and a learnable bias generated by spiking input from the MSNs, which were also modeled as integrate-and-fire neurons. In the critic component, VTA neurons represent the difference between the reward signal (*R*) from 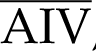 and the reward prediction signal (*RP*) from 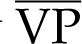 to generate the reward prediction error (*RPE*):

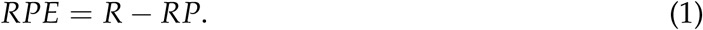

The RPE signal is transmitted to, and modulates plasticity at, the 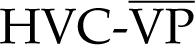 synapse to update the reward prediction, as well as the HVC-MSN synapse to drive actor learning.

To demonstrate the learning mechanisms and plasticity rules of the neural circuit model, we first consider a simple learning scenario at a single time in the song (a single note) and two motor output channels, one that improves song performance and one that impairs performance (Figure 2A). The actor component thus has one HVC neuron, and one LMAN neuron and MSN for each channel. Spiking of one LMAN neuron changes the song features (e.g., pitch) in a direction that improves the song performance (Figure 2A, blue) and spiking of the other LMAN neuron changes the song features in the opposite direction, impairing song performance (Figure 2A, red). Each HVC-MSN synaptic weight controls the MSN spiking activity, which then modulates the spiking probability of the respective LMAN neuron (Figure 2B). When the HVC-MSN weight is small, the LMAN neuron spikes at the baseline exploration probability. When the HVC-MSN weight is sufficiently large, HVC input activates the MSN to spike and consequently excites the downstream LMAN neuron to increase its spiking probability.

**Figure 2:**
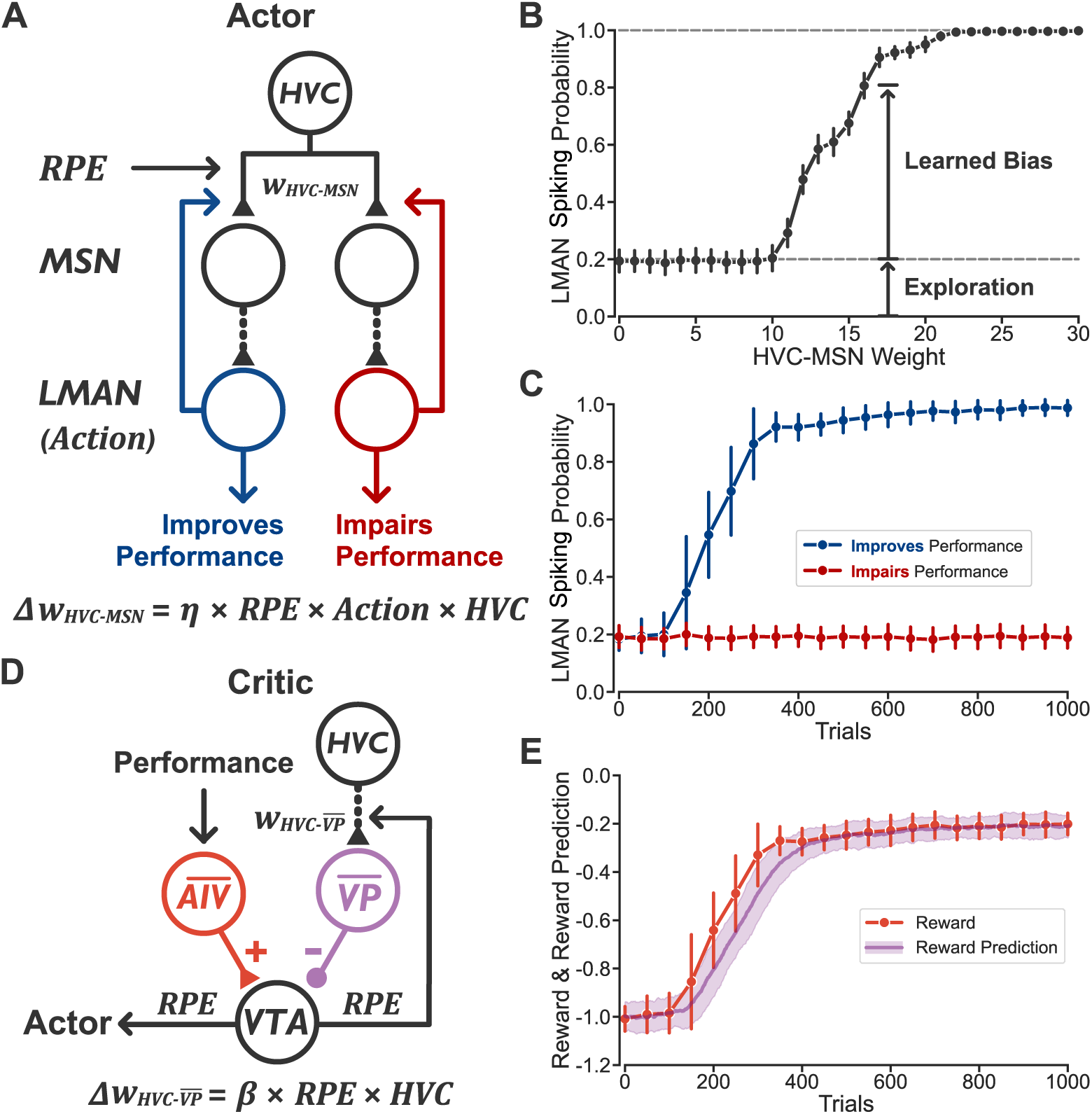
Actor-critic neural circuit model of song learning in a simple learning scenario. **(A)** Schematic illustration of the actor component. **(B)** LMAN spiking probability as a function of the HVC-MSN weight. **(C)** Changes in LMAN spiking probabilities across learning. **(D)** Schematic illustration of the critic component. **(E)** Expectation values of reward (orange, calculated from the firing probabilities in (C)) and the reward prediction (purple) throughout learning. Error bars in (C) and (E) and shaded band in (E) indicate standard deviation across 100 repetitions.

We implement the following actor learning rule for the HVC-MSN synapse suggested previously [27]:

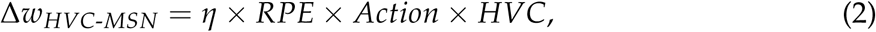

where *η* is a constant learning rate. The *Action* term represents the spiking of the LMAN neuron at each note and is modeled as a binary variable: *Action* = 1 if the LMAN neuron spikes at the corresponding note and *Action* = 0 if not. We refer to this actor learning rule as the ‘RPE learning rule’. Importantly, this plasticity rule requires the convergence of both temporal context signals (from HVC) and action signals (from LMAN). Anatomical evidence for such convergence and a detailed biophysical implementation of such a learning mechanism can be found in [62]. Physiological evidence for this learning mechanism can be found in [28]. We propose that the convergence of HVC and LMAN activity leads to an eligibility trace for synaptic plasticity [46, 63, 62], which maintains a memory of the HVC-LMAN coincidence until the arrival of the RPE signal from the evaluation pathway. Note that there is a time delay between the action produced by the LMAN neuron and the RPE signal produced after the song has been evaluated. We do not explicitly model this delay, under the assumption that the eligibility trace has to match such a delay for proper association between an action and the outcome to which it leads (Methods) [64, 65].

During learning, spiking of the performance-improving LMAN neuron leads to positive RPE signals and the corresponding HVC-MSN weight increases. Consequently, its spiking probability increases across learning (Figure 2C, blue). By contrast, spiking of the performance-impairing LMAN neuron leads to negative RPE signals, driving the corresponding HVC-MSN weight to decrease. Consequently, the HVC input to the MSN remains subthreshold and the LMAN spiking probability remains at the exploratory baseline (Figure 2C, red).

The RPE signal required for learning is generated by the critic component (Figure 2D). For the critic to learn the reward prediction, the HVC-VP synapse follows the critic learning rule:

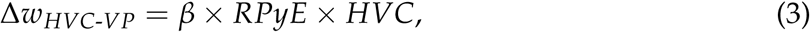

where *β* is a constant update speed. The HVC input can be considered to be binary, since each HVC neuron either exhibits a stereotyped bursting activity at its corresponding note or remains silent at the other notes. Thus, the HVC term can be assigned as a constant value of 1 when considering a single note. If the reward prediction is smaller than the actual reward received, the reward prediction error signal will be positive. This strengthens the 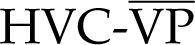 synapse and thus increases the reward prediction. As a result, the 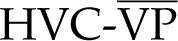 synaptic weight represents the reward prediction, and the critic learning rule performs the standard update of reward prediction:

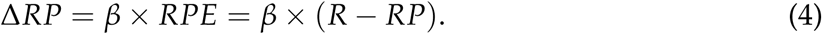

The above equation represents a geometric decay of the reward prediction toward the reward, with decay rate governed by the update speed *β*. During learning, the expected reward increases as the song performance improves (Figure 2E, orange), and the reward prediction tracks the expected reward with a lag caused by the finite update speed (Figure 2E, purple).

### 2.2 Song learning at each note as a two-armed bandit problem

The model described above has several interesting parallels to classical reinforcement learning theories. We first demonstrate that song learning at each note can be mapped to a canonical, well-studied problem in reinforcement learning known as the two-armed bandit problem. In this problem, an actor makes a binary choice between the left and right arms of a two-armed slot machine (Figure 3A). Each choice leads to a reward drawn from left- or right-arm distributions with different expectation values. Based on the chosen action and the reward received, the actor learns to prefer the action corresponding to the higher expected reward. Similarly, during song learning, at each note of the song, an LMAN neuron can be considered to be a two-armed slot machine, where the presence or absence of an LMAN spike corresponds to the choice of the right or the left arm. Each choice leads to a probabilistic reward with different expectation values depending on whether the generated song variation improves or impairs the song performance. Plasticity at the HVC-MSN synapse adjusts the spiking probability of the LMAN neuron based on the reward received. The RPE learning rule will, on average, strengthen the HVC-MSN synapse to increase the LMAN spiking probability if the LMAN spiking leads to a higher expected reward than the LMAN neuron not spiking (Figure 3B):

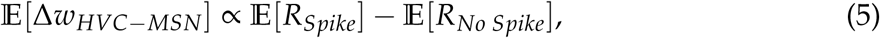

where E[*R_Spike_*] and E[*R_No Spike_*] represent the expected reward received when LMAN spikes or does not spike, respectively.

**Figure 3:**
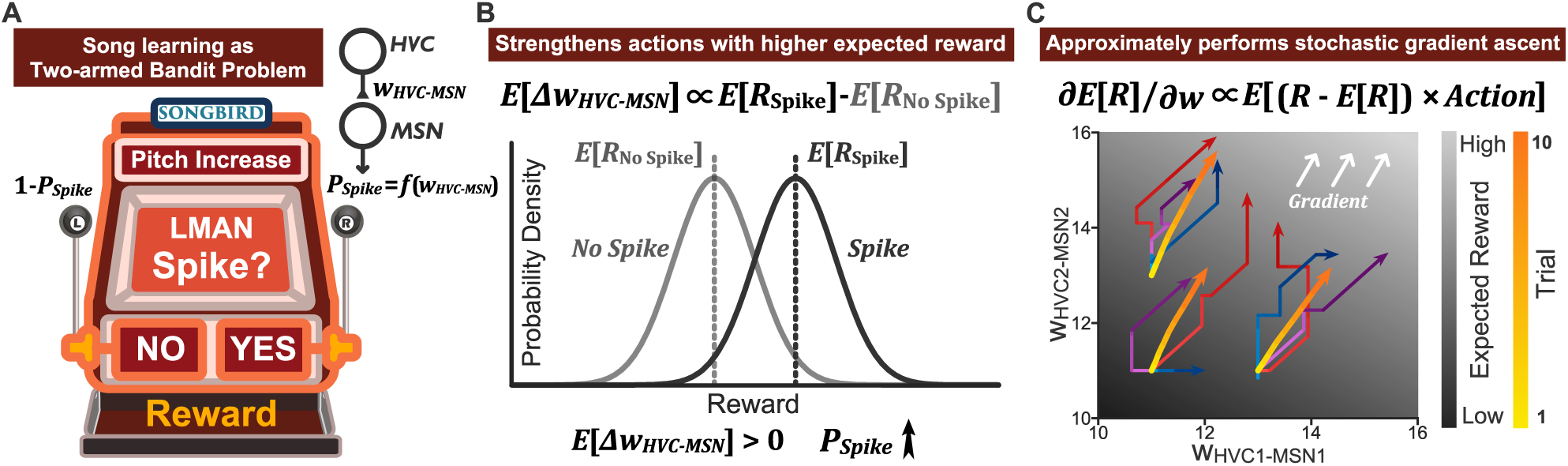
Song learning at one note is equivalent to a two-armed bandit problem. **(A)** The two-armed bandit problem. Each LMAN neuron (illustrated for one that increases the pitch) is analogous to a two-armed slot machine. Whether there is an LMAN spike or not corresponds to the choice of the right or left arm of the slot machine. The LMAN spiking probability (i.e., the probability of choosing the right arm) is a function of the HVC-MSN synaptic weight (Figure 2B). **(B)** If LMAN spiking leads to a higher expected reward than LMAN not spiking, the expected change of the HVC-MSN synaptic weight is positive, increasing the LMAN spiking probability. **(C)** The RPE learning rule approximately performs stochastic gradient ascent. Blue, red, and purple traces show three example learning trajectories of the two HVC-MSN weights starting from three different initial conditions. Yellow traces show the average learning trajectories across 1,000 repetitions, which approximately follow the direction of the gradient (white arrows). Trace colors from light to dark indicate trial 1 to 10.

We next show that the RPE learning rule for a single note is an approximate biological realization of the stochastic gradient ascent algorithm in reinforcement learning. Since the goal of reinforcement learning is to maximize the expected reward received, one class of algorithms seeks to change the learnable parameters in the direction of the gradient of expected reward, i.e., the direction along which the expected reward increases at the fastest rate. In the neural circuit model, the expectation value of the change of the HVC-MSN synaptic weights follows the direction of the gradient if the LMAN spiking probability follows a sigmoidal function of the weight (see Supplemental Information for proof). Mathematically:

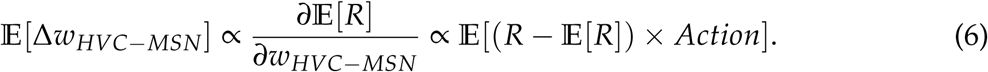

To demonstrate this gradient-following behavior, we performed simulations of a learning scenario in which there are two LMAN neurons that both improve the song performance but to different extents (Methods). The HVC-MSN weights evolve stochastically during learning, but on average approximately follow the direction of the gradient (Figure 3C). We note that this direct use of reward prediction error to move up the gradient of the expected reward contrasts with traditional gradient-based reinforcement learning algorithms [4] in which reward predictions appear only as an optional baseline parameter that is not involved in the calculation of the gradient (see Discussion). Thus, our results show how the widely observed reward prediction errors carried by dopaminergic neurons implement a biological stochastic gradient ascent algorithm, which, to our knowledge, has not been formally demonstrated. We also find that our results apply not only to the simple two-armed bandit problem but also to general reinforcement learning problems with multiple action choices and multiple notes (Supplemental Information).

In contrast to the previous scenario, where we only considered a single note, natural bird song is a sequential behavior spanning many notes controlled by HVC neurons firing at different time points. Thus, if each note in the song is independent of the other notes, then song learning can be considered to be a temporal sequence of independent two-armed bandit problems. However, songbird dopamine signaling is not temporally precise at the scale of a note; that is, the action mediated by an LMAN neuron at one note can lead to a dopamine signal that extends across multiple notes (several tens to a hundred milliseconds) [55, 56]. This temporal smearing of the dopamine signal leads to the question of whether there is interference between the learning of different notes. We thus tested whether a temporal sequence of actions can be accurately learned in the presence of such temporally imprecise dopamine signaling. We consider a pitch-increasing LMAN neuron learning at ten consecutive notes (representing a typical song syllable) by expanding the neural circuit model to have ten HVC-MSN pairs that converge onto the same LMAN neuron (Figure 4A). The target pitch is assumed to be higher than the baseline pitch only at the third note. Thus, the LMAN neuron needs to learn to increase its spiking probability precisely at the third note but not the others. Despite the temporally smeared dopamine signal, the LMAN neuron successfully learned the task (Figure 4B, 4C). Before learning, the LMAN neuron generates a sequence of random binary outputs across the ten notes. After learning, the LMAN neuron systematically fires at the third note but not the other notes (Figure 4B, 4C). For all notes, the critic generates reward predictions that track the actual expected rewards (Figure 4D). Despite the temporal imprecision of the dopamine signaling (Figure 4E, top), the model learns the correct behavior at millisecond precision, as the pitch trajectory shows an increase in pitch constrained within the target note (Figure 4E, bottom), much more precise than the timescale of dopamine signaling.

**Figure 4:**
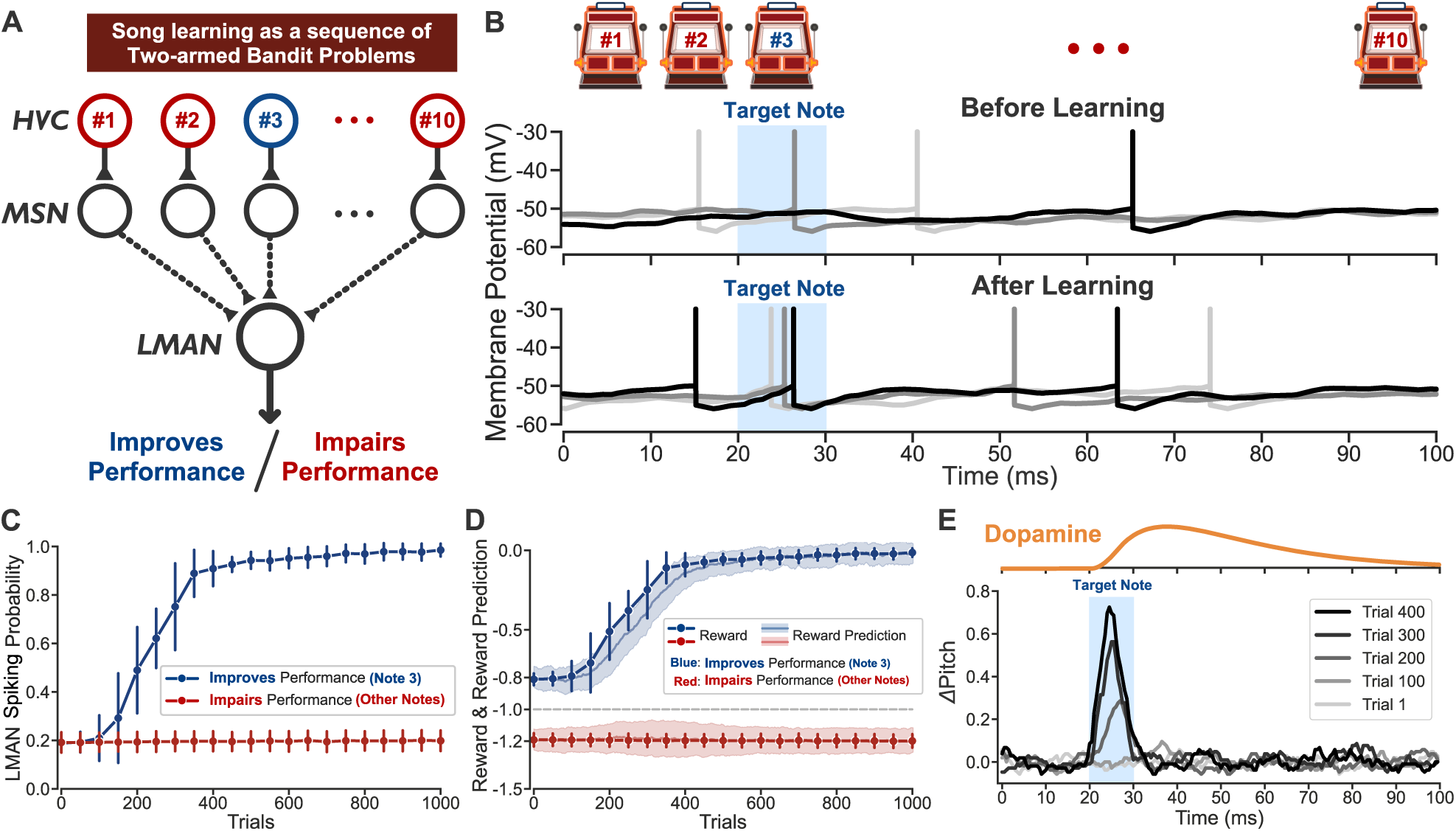
Temporally precise learning with imprecise reward in a sequence of two-armed bandit problems. **(A)** Learning task with ten consecutive notes. Spiking of a pitch-increasing LMAN neuron will improve or impair the song performance at a given note depending on whether the target pitch at that note is higher than the original pitch (note 3) or not (all other notes). **(B)** Three examples (light gray, dark gray, black traces) of the membrane potential of the LMAN neuron before learning (trial 1) and after learning (trial 1,000). Spikes are truncated at −30 *mV* for illustration. **(C)** LMAN firing probabilities throughout learning at note 3 (blue) and other notes (red). Error bars indicate standard deviation across 100 repetitions. (D) Expectation value of the reward and the reward prediction across learning. Error bars and shaded bands in (C) and (D) indicate standard deviation across 100 repetitions. **(E)** Top: dopamine signals (represented by VTA activity, see Methods) when the LMAN neuron fires at the third note of trial 1, averaged across 100 repetitions. Bottom: pitch trajectories (Methods) after different numbers of learning trials, averaged across 100 repetitions, then plotted relative to the average pitch of trial 1.

### 2.3 Model performs precise song learning that matches behavior

In this section, we demonstrate that the neural circuit model achieves precise song learning that matches behavior in real song learning experiments. We consider the learning of multiple notes in the context of a conditional auditory feedback (CAF) paradigm [66, 38] in which disruptive auditory feedback (typically a broadband noise) is selectively played in real time to the singing bird, simulating a large vocal error. The disruptive feedback is made contingent on the pitch of a target syllable of the song by providing this feedback whenever the pitch crosses a certain threshold during the target syllable (Figure 5A). In songbirds, such a manipulation causes pitch learning in a direction where the disruptive feedback is avoided. For example, playing disruptive feedback whenever the pitch is above the threshold causes the average pitch of the target syllable to decrease until the distribution of the average pitch is mostly below the threshold (Figure 5B).

**Figure 5:**
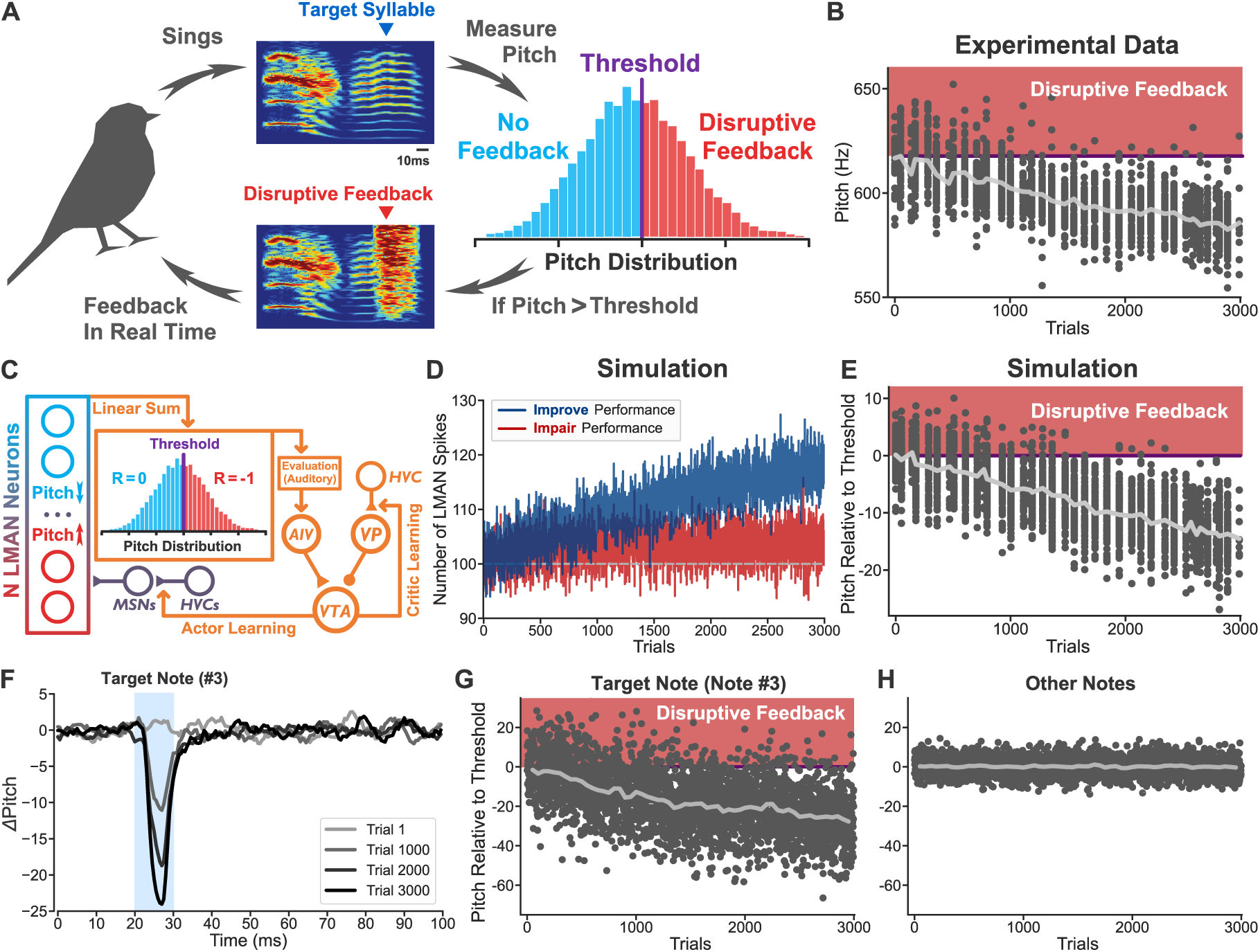
Model performs temporally precise song learning and matches experimental results. **(A)** Illustration of the conditional auditory feedback (CAF) experimental paradigm. Before learning, the distribution (across trials) of the average pitch (across time) of a target syllable is measured. During learning, a disruptive auditory feedback is provided to the singing bird whenever the pitch within the target syllable goes higher than a threshold given by the mean value of the average pitch distribution. **(B)** An example trajectory of the average pitch of the target syllable during one day’s learning (first 3,000 trials shown). Each dot represents one trial and trials are grouped together into bins along the x-axis, with trials close in time aligned vertically (Methods). The light gray line shows the mean of each bin. **(C)** Architecture of the neural circuit model with *N* = 1, 000 LMAN neurons, half of which decrease the pitch (blue, improve performance) and half of which increase the pitch (red, impair performance). Each LMAN neuron receives input from and sends feedback to 10 MSNs, each of which is driven by an HVC neuron representing 1 of 10 consecutive notes (Figure 4A). For each note, the pitch in the model is defined as the difference in the number of spikes generated by the two groups, and the reward is defined to be -1 if the pitch is higher than the threshold and 0 if the pitch is lower than the threshold. **(D-E)** Simulation of 3,000 trials of the CAF experiment. **(D)** Number of spikes generated by the performance-improving group (blue) and the performance-impairing group (red) throughout learning. Horizontal gray dashed line indicates the baseline number of spikes in expectation, corresponding to the baseline LMAN spiking probability. **(E)** Average pitch of the target syllable throughout learning. Trials are binned as in panel B. **(F-H)** Simulation of 3,000 trials in which the CAF was targeted exclusively to a single note [61]. **(F)** The average pitch trajectory (across 100 repetitions) after different numbers of trials in which only the third note is targeted, plotted relative to the average pitch of trial 1. **(G)** Average pitch of the targeted note throughout learning. **(H)** Average pitch of the non-targeted notes throughout learning, averaged across the non-targeted notes. Histograms in (A) and data in (B) are adapted from [38].

To model these experimental data, we expanded the neural circuit model from the previous section to 1,000 LMAN neurons, to match the order of magnitude of the number of LMAN neurons that control one feature of the song (Figure 5C). The LMAN neurons were divided evenly into two groups. The spikes generated by one group decrease the pitch (improve performance) and those generated by the other group increase the pitch (impair performance). As seen in Figure 5D, the number of spikes generated by the performance-improving LMAN neurons increases across learning and the number of spikes generated by the performance-impairing LMAN neurons remains near the baseline. As a result, the pitch distribution shifts below the threshold (Figure 5E), similar to the experimental results.

We next modeled a variation of the previous experiment, in which the disruptive feedback is made contingent on a single note rather than an entire syllable of the song. Experimentally, songbirds can achieve temporally precise learning and only change the pitch at the target note [61]. Similarly, our neural circuit model learns to only change the pitch of the target note to avoid disruptive feedback, while the pitches of the other notes remain at baseline (Figure 5F-H).

### 2.4 Fast update speed of reward prediction is required to avoid the temporal interference problem

The results above demonstrate that temporally precise learning can be achieved with temporally imprecise dopamine signaling. Here we show the condition for this to occur, and how learning at different notes can interfere with one another when this condition is not met.

We first show that a temporally smeared dopamine signal can lead to temporal interference between learning at neighboring notes if the update speed of reward prediction is slow. If the dopamine signaling were temporally precise, the action at a single note would elicit a dopamine signal that is temporally localized to that note (Figure 6A, top). However, dopamine signaling in the songbird brain is smeared across time so that an RPE at one note elicits a dopamine signal that spans multiple notes [55, 56] (Figure 6A, bottom). To illustrate the interference problem that can be caused by such temporal imprecision, we consider a scenario in which a single LMAN neuron learns at five consecutive notes. In this scenario, we set the target pitches at each note so that LMAN spiking improves performance at the first four notes (Figure 6B, blue) but impairs performance at the fifth note (Figure 6B, red). Ideally, when the LMAN neuron spikes at note 5, the dopamine signal should be negative at note 5 to reflect that the neuron’s spiking generated an error. In contrast, when the LMAN neuron spikes at notes 1-4, the dopamine signal should be positive to drive plasticity that increases spiking at these times. However, when the dopamine response is smeared out, spiking of the LMAN neuron at notes 1-4 produces a long tail in the RPE that can bleed into note 5, producing a positive dopamine signal that can drive erroneous learning at note 5. If the LMAN spiking is sparse, as typically occurs early in learning (Figure 6C, left), then the RPE tail will not occur frequently enough to cause erroneous learning. However, after the LMAN neuron has begun to learn to spike at notes 1-4, at the middle stage of learning (Figure 6C, middle), the LMAN spiking at notes 1-4 occurs more densely. This can lead to a consistently occurring RPE tail that is large enough to cause the dopamine signal at note 5 to be always positive, leading to erroneous learning at this note. Later in learning (Figure 6C, right), the reward prediction eventually catches up so that the improved performance at notes 1-4 is properly accounted for, and the dopamine signal at note 5 reflects only the error occurring at this note. Overall, these toy examples illustrate how interference from previous notes can lead to incorrect, positive RPE signals (Figure 6D) and an associated erroneous change in LMAN spiking probability (Figure 6E).

**Figure 6:**
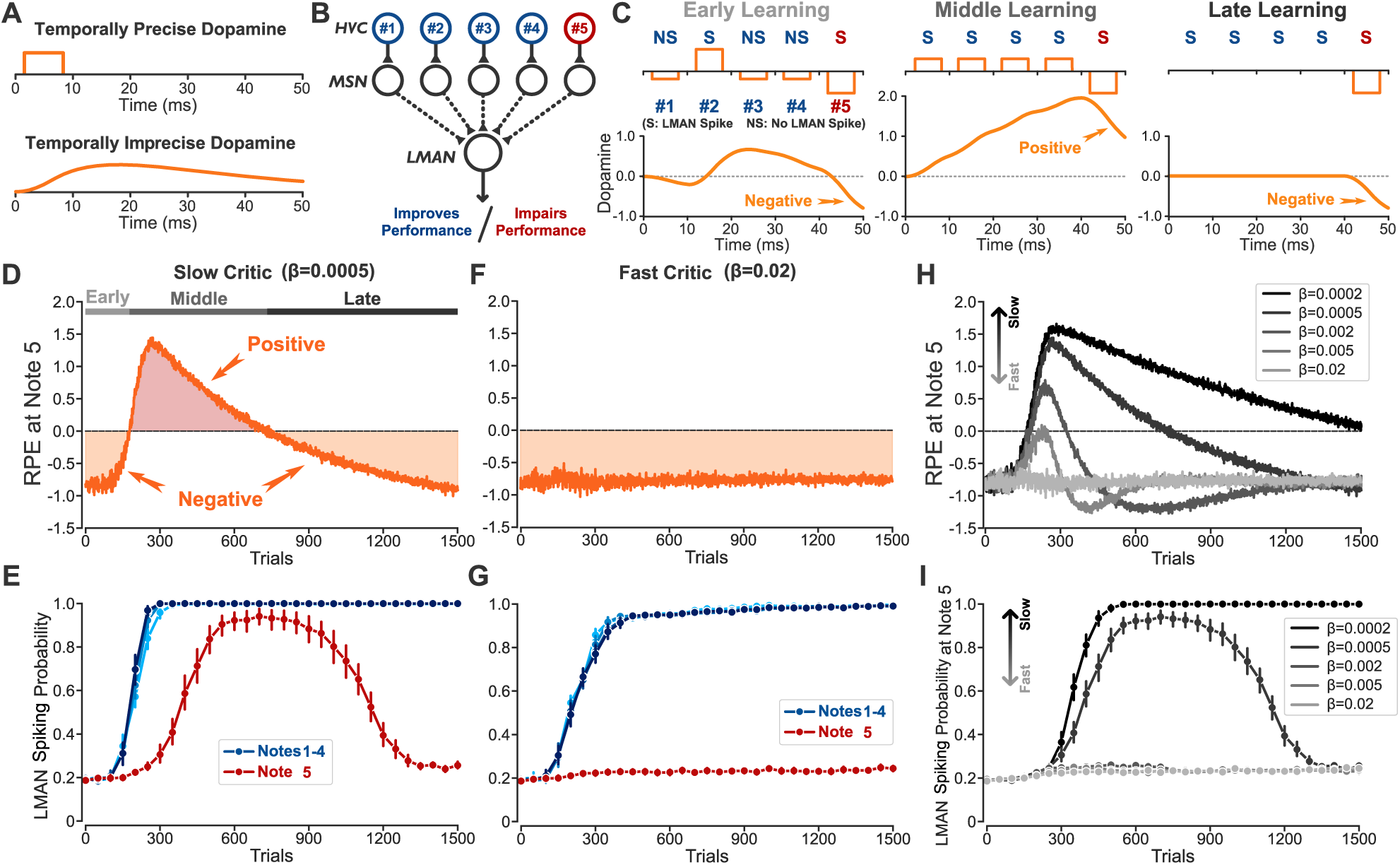
Temporal interference causes incorrect learning that can be resolved by a fast update speed of reward prediction. **(A)** A temporally precise dopamine signal should be constrained to the note causing the RPE (top). A temporally imprecise dopamine signal extends to future notes (bottom). **(B)** Diagram of a learning task with five consecutive notes in which the LMAN neuron is assumed to improve the performance at notes 1 - 4 but impair the performance at note 5. Blue: notes at which LMAN spiking improves performance. Red: note at which LMAN spiking impairs performance. **(C)** Representative examples of the dopamine signals (orange) during the early, middle, and late stages of learning when dopamine is temporally precise (top) or when it is temporally imprecise (bottom). The shown sequences of LMAN spiking (S) and LMAN non-spiking (NS) are chosen to highlight the dopamine signaling in cases in which the LMAN neuron fires at note 5 and thereby impairs performance. **(D and E)** Learning results with a slow critic. D: The RPE signal at note 5 (averaged across 100 repetitions) throughout the early, middle, and late stages of learning. Here, the RPE signal is defined as the dopamine value at the end of the note (see Methods). E: LMAN firing probabilities at notes 1 - 4 (blue, lighter colors represent earlier notes) and note 5 (red) throughout learning. **(F and G)** Same as (D) and (E) but with a relatively fast critic. (H) The RPE signal at note 5 (averaged across 100 repetitions) for different values of *β*. **(I)** The LMAN firing probability at note 5 for different values of *β*. All error bars indicate 95% confidence intervals across 100 repetitions. Note that we use 95% confidence intervals here, instead of standard deviation as in other figures, because the standard deviation range may include firing probabilities that exceed 1.

The extent to which this smearing in the dopamine signal causes erroneous learning depends on whether the update speed of reward prediction is sufficiently fast. For example, the *β* value used in previous sections (*β* = 0.02) leads to a correct, negative dopamine signal throughout learning at note 5 (Figure 6F), so that there is no incorrect learning (Figure 6G). To further examine how the update speed of reward prediction affects erroneous learning, we simulated five different update speeds of reward prediction. These simulations show that the temporal interference and incorrect learning problems are resolved if the critic is sufficiently fast (Figure 6H,I).

A slow critic also causes oscillations in the LMAN firing probabilities at notes 1-4. Specifically, our simulations demonstrate that, following the initial correctly learned increases, the firing probabilities then partially unlearn before relearning over a longer timescale (Figure S1). We discuss the mechanistic origin of this problem in the Supplemental Information.

### 2.5 Fast update speed of reward prediction is required to avoid interference across motor channels

In the above section, we showed that a slow update speed of reward prediction can lead to interference across time using the example of a single LMAN neuron learning at multiple notes. Here, we show a similar interference effect when there are many LMAN neurons controlling different motor channels that contribute to song learning (e.g., one group of LMAN neurons that increases the pitch and improves song performance and another that decreases the pitch and impairs song performance). Dopamine signaling is not only temporally imprecise and smeared across time (Figure 6A), but also ‘spatially’ imprecise and smeared across motor channels (Figure 7A). In the actor-critic framework, it would be ideal if the actors (i.e., the LMAN neurons that control different motor channels) were evaluated by independent critics such that the actors do not interfere with each other (Figure 7A, top). However, experimental observations [55, 56] suggest that the song system likely contains a single critic representing the overall song performance (Figure 7A, bottom). Having a single critic can cause interference when learning in performance-improving actors leads to a positive RPE that consequently causes incorrect learning in the performance-impairing actors. This is similar to the temporal interference problem treated in the previous section, in which learning of earlier notes causes a temporally imprecise positive RPE and incorrect learning of later notes (Figure 6).

**Figure 7:**
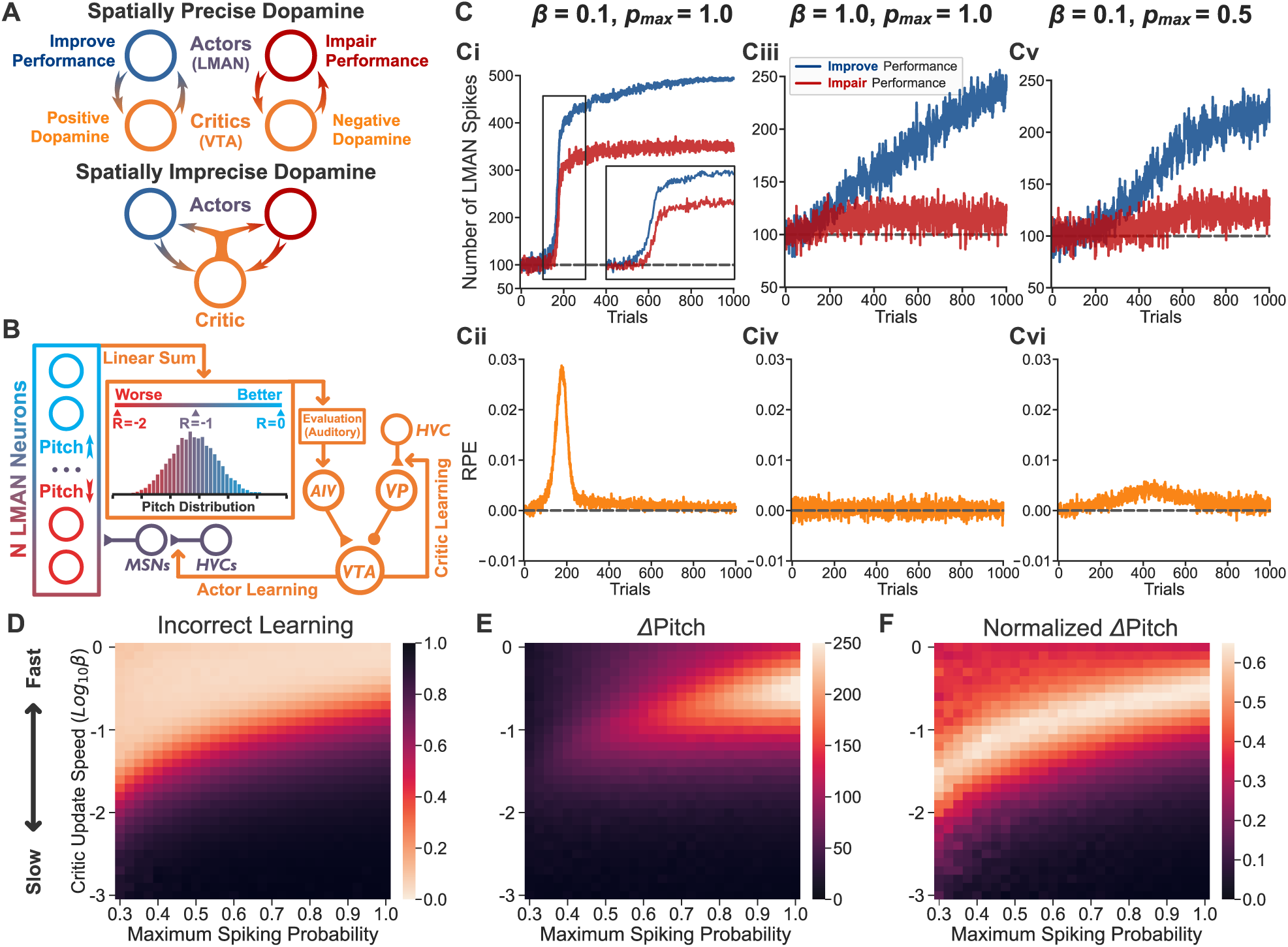
Spatial interference causes incorrect learning that can be resolved by a fast critic. **(A)** Top: Spatially precise dopamine signaling in which each of two motor channels (actors) is evaluated by its own corresponding critic so that there is no spatial interference across actors. Bottom: Spatially imprecise dopamine signaling in which a global signal evaluates and guides the learning of both motor channels simultaneously. **(B)** Diagram illustrating a pitch-increase learning task with *N* = 1, 000 LMAN neurons, half of which increase the pitch and half of which decrease the pitch. The reward is assumed to be proportional to the pitch, which is defined as the difference between the number of spikes generated by the pitch-increasing group and the pitch-decreasing group, normalized between -2 and 0. **(C)** Total LMAN population firing and RPE across learning for three different sets of update speeds *β* and maximum LMAN firing probabilities *p_max_*. Top (i, iii, v): example trajectories of the total spike counts in the pitch-increasing (blue, improve performance) and pitch-decreasing (red, impair performance) groups. Inset in i is a zoomed-in view from trial 100 to 300. Bottom (ii, iv, vi): RPE averaged across 1,000 repetitions. Horizontal dashed lines indicate the baselines. **(D)** The extent of incorrect learning (defined as the spike count increase from baseline in the performance-impairing group, normalized by the maximum possible increase) for different values of *β* and *p_max_*. Learning was measured at trial 1,000 and averaged across 100 repetitions. **(E)** Same as (D), but showing the amount of learned pitch increase. **(F)** Same as (E), but normalized by the maximum possible pitch increase. Note that the color bar in (D) is reversed from (E, F) for consistency; lighter colors represent better results.

To investigate this issue, we consider song learning with multiple LMAN neurons controlling the pitch of a single note. We simulate learning with 1,000 LMAN neurons, half of which increase the pitch and half of which decrease the pitch. Unlike the conditional auditory feedback experiment in which the reward is a binary function (Figure 5C), for the case of natural feedback in which song performance is evaluated by comparison with a stored template [53], we assume that the reward is linear with respect to song features. Consequently, the closer the pitch is to the target pitch, the higher the reward will be (Figure 7B). If the baseline pitch starts below the target pitch, the goal of learning is to increase the firing probabilities of the pitch-increasing (i.e., performance-improving) LMAN neurons but not the pitch-decreasing (i.e., performance-impairing) LMAN neurons.

Similar to the temporal interference problem described in the previous section, spatial interference across motor channels also causes incorrect learning if the critic is not fast enough. For example, with an update speed of *β* = 0.1, corresponding to a 10-trial time constant of learning, we found that there was an abrupt, erroneous increase in spiking of the performance-impairing LMAN neurons (Figure 7Ci). The cause of this problem is a positive feedback loop between LMAN spiking and reward, which can lead to excessively positive RPE signals (Figure 7Cii). Specifically, the performance-improving group first learns to produce more spikes, leading to an increase of reward and positive RPE signals, which in turn correctly drives the performance-improving group to produce even more spikes. However, the excessively positive RPE signal caused by this positive feedback loop also incorrectly drives learning in the performance-impairing group, resulting in non-optimal performance. By contrast, with a very fast critic (*β* = 1), the reward prediction can keep up with the increasing reward, moderating the RPE signals to an appropriately small size so as to avoid triggering the feedback loop (Figure 7Civ). In this case, only the performance-improving group learns to produce more spikes, while the performance-impairing group remains close to the baseline level (Figure 7Ciii).

Increasing the reward prediction update speed is not the only way to tame the positive feedback loop leading to excessively positive RPE signals. One alternative is to reduce the learning rate of the actor, i.e., the rate of plasticity at the HVC-MSN synapse. A slower actor learning rate would have a less stringent requirement on the speed of critic learning but would inevitably make learning less efficient by requiring more trials to learn (Figure S2). In this study, we fix the actor learning rate either to match experimental results (Figure 5) or to achieve learning in the model on the 1000-trial timescale of a day of natural birdsong learning [67]. Another possibility to mitigate the excessively positive RPE signals is to set an upper bound on the maximum spiking probability of the LMAN neurons, *p_max_*. This reduces the maximum amount of learning that can be achieved in a day (Figure 7Cv), and thus the maximum amount of reward. Consequently, the positive reward prediction error is reduced despite the fact that the reward prediction still lags behind the increase of reward (Figure 7Cvi). These alternative approaches can also mitigate the temporal interference problem.

The spatial interference described above not only causes incorrect learning but also impacts the overall rate of learning, i.e., the rate at which song errors are corrected. To quantify this, we systematically examined the extent of incorrect learning in the performance-impairing group of neurons (Figure 7D) and the overall amount of correct pitch change (ΔPitch) (Figure 7E,F) under different critic update speeds and maximum LMAN spiking probabilities. Learning with a sufficiently fast update speed avoids incorrect learning regardless of the maximum spiking probability, whereas a slower update speed requires a smaller maximum spiking probability (Figure 7D). The pitch is learned most quickly when there is no limit on the LMAN spiking probability, i.e., *p_max_*= 1, and when the update speed is approximately *β* = 0.3 (Figure 7E), close to that measured experimentally [53] (see Discussion). This ideal update speed is slightly slower than the maximum update speed of *β* = 1, leading to a small amount of positive feedback between the RPE and LMAN spiking, as described above. This can boost the overall learning rate without causing notable incorrect learning in the performance-impairing neurons. Notably, comparable fast updating has been observed in animal studies across species (see Discussion). To better isolate the effect of the reward prediction update speed, we computed the amount of pitch increase normalized by the maximum amount achievable for a given *p_max_*. Again, the best normalized performance is obtained when the update speed is fast enough to restrain the incorrect learning (Figure 7F).

The positive RPE signals that cause the incorrect learning problem scale with the number of LMAN neurons that drive the increased reward (Supplemental Information). As a result, a much faster reward prediction update speed is required to avoid incorrect learning when the number of LMAN neurons is large (Figure S3).

## 3 Discussion

We built a spike-based actor-critic neural circuit model of the songbird vocal learning system with biologically plausible learning rules. The model shows that song learning can be understood as a sequence of two-armed bandit problems, in which the two choices correspond to whether LMAN neurons spike or remain silent at specific moments in the song. RPE-based synaptic plasticity in this circuit approximately implements stochastic gradient ascent on song performance, allowing the model to learn accurately and reproduce behavioral effects observed in songbirds subjected to targeted song disruptions. By modeling both the actor and critic components of the circuit, we identified two forms of interference that may be broadly relevant to RPE-driven learning: temporal interference across actions occurring close in time and spatial interference across neurons controlling different motor channels. Both forms of interference arise when the reward prediction is updated too slowly relative to improvements in performance, producing excessively positive RPEs that can reinforce neurons that were not responsible for the improvements. Our results therefore predict that a fast critic is required for accurate and efficient reinforcement learning when global neuromodulatory signals provide temporally and spatially imprecise feedback.

### Implications for reinforcement learning circuitry in other biological systems

Reinforcement learning across the animal kingdom requires the convergence of three pieces of information: context, action, and performance evaluation. In the songbird neural circuit model, this convergence occurs at the striatal medium spiny neurons (MSNs). First, the HVC neurons fire robustly in sequence and provide a temporal context for the song. Second, the LMAN neurons generate exploratory actions and the feedback connections from LMAN neurons to MSNs serve as efference copies of the chosen actions. Third, VTA dopaminergic neurons project to MSNs and signal the reward prediction error that represents an evaluation of the behavioral performance. While HVC and LMAN are specialized brain areas in songbirds, a temporal context signal such as that provided by the HVC neurons should be important for reinforcement learning of sequential behaviors more broadly. Similarly, an efference copy of action is ubiquitous across the animal kingdom [68] and could be a crucial piece of information for various learning systems [69, 70].

The anatomy of mammalian cortico-basal ganglia circuitry may share a similar functional organization to that of the birdsong learning system. Similar to the HVC and LMAN convergence onto MSNs in songbirds, mammalian MSNs receive their main cortical inputs from intratelencephalic (IT) and pyramidal tract (PT) neurons, and the striatum is unique among all subcortical regions in receiving both IT and PT inputs [71]. IT and PT inputs play different functional roles, with the IT neurons conveying sensory information and the PT neurons in the motor cortex conveying an efference copy of motor commands [72]. We interpret such sensory information as a context signal, analogous to the timing information conveyed by HVC. Anatomically, the HVC and LMAN neurons in songbirds converge onto the MSNs: the HVC synapses target spine heads and the LMAN synapses preferentially target dendritic shafts [62]. Thus, the songbird circuitry suggests that examining the anatomical properties (i.e., targeting the spine or shaft) of the inputs onto MSNs can help to clarify their functional purpose in motor production and learning. Mammalian MSNs receive cortical and thalamic inputs onto both the spines and shafts, with cortical inputs predominantly onto the spines and thalamic inputs relatively more abundant onto the shafts [73, 74, 75, 76]. Interestingly, PT neurons project onto the thalamus [77] including the Pf region that projects heavily to the striatum [78, 79, 80]. If the PT-recipient thalamic neurons project onto the dendritic shafts of striatal MSNs, this would suggest a functional analogy with the birdsong system, in which the PT-driven thalamic activity provides an efference copy of motor actions to the striatum. More broadly, we hypothesize that functional and anatomical convergence of context and action similar to that in the songbird circuitry could be a general principle for biological reinforcement learning systems [70].

#### Implications for the function of reward prediction error in reinforcement learning

Our modeling suggests that the reward prediction errors observed during songbird learning can support a policy-based reinforcement learning algorithm. Traditional reinforcement learning algorithms primarily fall into two categories: value-based and policy-based [4]. Value-based reinforcement learning involves explicitly learning the values (i.e., expected rewards) of different states or actions, with these values used to decide which action to take. These values are updated by reward prediction errors, and the observation that midbrain dopaminergic neurons signal RPEs has therefore been widely interpreted as evidence for a value-based learning mechanism in the brain [17, 18, 81, 82, 12, 5]. By contrast, policy-based reinforcement learning can optimize a parameterized behavioral policy directly, rather than deriving choices from learned values of alternative actions [83, 84, 85, 86]. Specifically, policy gradient methods learn to increase the reward through stochastic gradient ascent. Our neural circuit model of song learning demonstrates the feasibility of such a policy gradient mechanism, with the corticostriatal plasticity of HVC-MSN synapses directly adjusting the spiking probability of the downstream LMAN neuron without explicitly encoding action values. Formally, the classic theory underlying policy gradient methods only requires rewards, not reward prediction errors [4]. Here, we show that reward prediction errors can not only mediate value learning but can also implement stochastic gradient ascent (Supplemental Information). We note that classic theories often do include a reward prediction error term by subtracting the expected reward from the reward signals. However, this subtraction is not part of the policy gradient calculation. Instead, it is used to recenter the reward signals to reduce the variance of learning. By contrast, our theory demonstrates an alternative formulation of policy-based reinforcement learning in which the reward prediction error itself is the fundamental quantity underlying stochastic gradient ascent (Supplemental Information).

#### Fast critic as a general principle of biological reinforcement learning

A fast update speed of reward prediction has been reported during natural song learning in juvenile songbirds [53], indicating an update speed of about *β* = 0.3 (Methods), closely matching the predicted optimal update speed in our model (Figure 7E). Our results provide an explanation for such a fast critic and suggest that it may be a general principle of biological reinforcement learning. Indeed, fast updating of reward prediction has been reported across species. In a saccade timing task performed by monkeys, it was found that an update speed of *β* = 0.7 best captures the firing properties of the midbrain dopaminergic neurons that encode reward prediction error [81]. In a two-choice saccade task performed by monkeys [87], dopaminergic neurons show an update speed *β* of about 0.5. Another study, in rats performing an instrumental nose-poke task [88], found that the update speed of reward prediction depends on the target region of the dopaminergic neurons, with a *β* value of about 0.1 for ventral striatum, about 0.2 for dorsomedial striatum, and as fast as 0.5 for dorsolateral striatum, also consistent with our prediction of a fast critic.

From the standpoint of optimizing critic accuracy alone, while ignoring the dynamics of the interactions with the actor, one would select an update speed that minimizes reward prediction error. Minimizing RPE entails a bias-variance tradeoff: faster update speeds reduce bias by allowing reward predictions to track changes in reward more closely, whereas slower update speeds reduce variance by averaging out trial-by-trial noise. Consequently, an intermediate update speed is optimal, and an idealized analysis (Supplemental Information) suggests a range of *β* = 0.01 − 0.1 for optimizing this bias-variance tradeoff. Thus, from the perspective of solely having an accurate critic, the fast update speeds observed in animal studies may seem counterintuitive. Here, we demonstrate that such an analysis critically neglects the dynamical interactions between the actor and critic. When these interactions are taken into account, a fast critic is essential to avoid interference problems that disrupt proper actor learning.

#### Consolidation of learning

In this study, we only considered the fast learning process in the songbird basal ganglia pathway (the learning pathway), which occurs on the order of a day. A slower learning process happens in the motor pathway, where the learned song variations generated by LMAN neurons transfer to the HVC-RA connections [38, 41], a process generally referred to as consolidation of learning. Following the consolidation of learning, the motor pathway alone is able to generate the learned variations and the learning pathway is reset to an unlearned state, ready for new learning processes. Mechanistically, consolidation can be readily carried out by heterosynaptic plasticity [89], in which the co-activation of HVC and LMAN neurons converging onto RA neurons leads to long-term potentiation of HVC-RA synapses. Concurrently, the HVC-MSN synaptic weights in the learning pathway gradually decay to values below the threshold for driving MSNs, possibly due to long-term depression resulting from co-activation of HVC neurons and MSNs without dopaminergic reinforcement. Including such a consolidation process would enable future models to describe the full process underlying birdsong learning, from the initial adjustments of song performance via motor exploration-driven changes in the basal ganglia-cortical circuitry to the final consolidation of learning within the direct cortical pathways.

## Methods

We built a neural circuit model of the song learning circuitry consisting of two interacting components (Figure 1B): the actor component (HVC, MSNs, and LMAN) and the critic component (song evaluation, HVC, 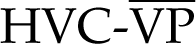, 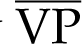, and VTA). The motor pathway (from HVC to RA and lower motor areas) is not explicitly modeled since its activity is stereotyped within the timescale of learning considered here and thus represents a behavioral baseline.

### Neural Circuit Model: Actor

The actor component consists of HVC, MSNs, and LMAN. The number of LMAN neurons, *N_LMAN_*, depends on the task. *N_LMAN_*= 2 in Figure 2, where we consider one pitch-increasing and one pitch-decreasing LMAN neuron to demonstrate how the model performs reinforcement learning. *N_LMAN_* = 2 in Figure 3 to show that the RPE learning rule approximately performs stochastic gradient ascent. *N_LMAN_* = 1 in Figures 4 and 6, where we focus on the temporal aspect of the learning and thus only consider a single LMAN neuron for simplicity. In Figures 5 and 7, where we examine realistic pitch learning tasks, *N_LMAN_*= 1, 000 to match the order of magnitude of the number of LMAN neurons that control one song feature (there are ∼10,000 LMAN neurons in the songbird brain [90, 91] controlling ∼10 vocal muscles [92] or song features [56]). For each LMAN neuron, one HVC–MSN pair controls one note of the song. Thus *N_HVC_* = *N_MSN_* = *N_LMAN_*× *N_note_*, where the number of notes *N_note_* = 1 in Figures 2, 3, and 7, *N_note_* = 10 in Figures 4 and 5, and *N_note_*= 5 in Figure 6.

Each note is considered to occupy a 10 *ms* window. Relative to the starting point of a note, the corresponding HVC neuron is characterized by a burst of four stereotyped spikes occurring at −2 *ms*, 0 *ms*, 2 *ms*, and 4 *ms*. These values account for the propagation delay from the HVC neuron to the MSN and LMAN neuron, such that a strengthened HVC-MSN connection after learning tends to activate the downstream LMAN neuron to fire within the corresponding note. The MSN and LMAN neurons are modeled with leaky integrate-and-fire neurons, with a resting potential *E_L_*, a firing threshold *V_thr_*, a reset potential *V_reset_*, a membrane capacitance *C_m_*, a leak conductance *g_L_*, and a refractory period *T_re f_* . For the MSN, *E_L_*= −80 *mV*, *V_thr_* = −50 *mV*, *V_reset_* = −55 *mV*, *C_m_*= 0.1 *nF*, *g_L_* = 20 *nS*, *T_re f_* = 1 *ms*. MSNs have a very hyperpolarized resting potential [93, 94]. For the LMAN neuron, *E_L_*= −65 *mV*, *V_thr_* = −50 *mV*, *V_reset_* = −53 *mV*, *C_m_*= 0.5 *nF*, *g_L_* = 25 *nS*, *T_re f_* = 1 *ms*. The LMAN neuron is modeled with a relatively high reset potential such that spiking probabilities across notes are more independent (otherwise, when the LMAN neuron learns to fire more at one note, the spiking probability at the next note will be reduced).

For the MSN and LMAN neurons, the subthreshold membrane potential, *V*(*t*), follows:

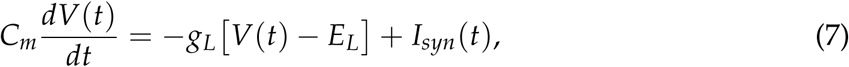

where *I_syn_*(*t*) represents the total synaptic current flowing into the cell.

The synapses onto MSNs receive excitatory HVC input and an excitatory external noisy input, both mediated by AMPA receptors. The noisy input is modeled as a Poisson spike train representing the sum of background inputs, with rate 1.3 *kHz*, which produces a subthreshold fluctuation of voltage. The total synaptic current is given by

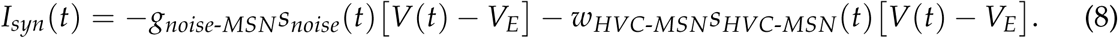

*V_E_* = 0 *mV* is the reversal potential of the AMPA receptors. *s*(*t*) describes the strength of presynaptic input (see below). *g_noise_*_-*MSN*_ = 1 is a constant synaptic weight of the noisy input and *w_HVC_*_-*MSN*_ is a learnable synaptic weight between the HVC neuron and MSN. Unless noted otherwise, *w_HVC_*_-*MSN*_ is initialized with a subthreshold value of 8 and constrained to have a lower bound of 0 and an upper bound of 30 (Figure 2B). In the conditional auditory feedback task (Figure 5), *w_HVC_*_-*MSN*_ is initialized to a value of 10 to better match the experimental data.

The LMAN neurons receive excitatory MSN input and an excitatory external Poisson input with rate 1.85 *kHz*, which produces a slightly suprathreshold input to elicit a baseline exploratory spiking probability of around 0.2 at each note, mimicking the spontaneous firing rate of the LMAN neurons during singing without MSN input [35, 95]. The total synaptic current onto an LMAN neuron is given by

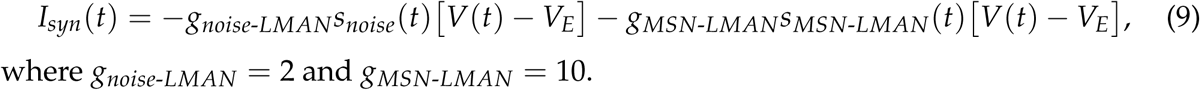

where g_noise-LMAN_ = 2 and g_MSN-LMAN_ = 10.

The strength of presynaptic input, *s*(*t*), is described by

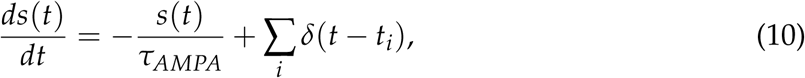

where *τ_AMPA_* = 2 *ms* and *t_i_*represents the time of presynaptic spikes. *δ* is a delta function such that *s*(*t*) increases by 1 when there is a presynaptic spike at time *t* = *t_i_*.

During learning, the spiking activity of each LMAN neuron at each note is characterized by a binary variable termed *Action*. If the LMAN neuron generates a spike within the corresponding 10 *ms* window, *Action* = 1; otherwise, *Action* = 0. *Action* of one LMAN neuron at each note is assumed to increase or decrease the pitch of that note (defined discretely for each note) by 1 unit. The continuous pitch trajectories are obtained by computing the LMAN firing rate (Figure 4E) or the difference between the number of LMAN spikes in the pitch-increasing and pitch-decreasing groups (Figure 5F) in a 5 *ms* sliding window in time steps of *dt* = 0.05 *ms*, and then normalized by the discrete pitch calculated from the *Action* at each note (Figures 4C and 5G).

### Neural Circuit Model: Critic

The critic component consists of HVC, 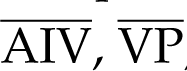, 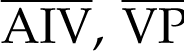, and VTA, with the activity of each brain area described by a firing rate *r*(*t*). Relative to the starting point of a note, the HVC input *r_HVC_*(*t*) received by 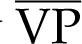 is characterized by a square wave of amplitude 1 from 2 *ms* to 8 *ms*. The evaluation signal *R*(*t*) received by 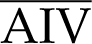 is characterized by a square wave with the same shape, and with amplitude that equals the value of the reward.

In songbirds, the timing of evaluation signals in the critic component of the circuit does not align perfectly with the timing of action-generating signals in the actor component. Instead, there is a delay from the time of generation of LMAN spikes to the song evaluation and the generation of VTA dopamine signals. We do not explicitly model this time delay between the actor and the critic signals, instead assuming that the eligibility trace has to match such a delay for proper association between the LMAN spike and the dopamine signal it leads to [64, 65]. In the figures, we align the RPE signal with the LMAN activity or song features to indicate a functional correspondence (Figures 4E, 6C). In our model, we assume for simplicity that the eligibility trace is temporally precise and that temporal imprecision arises only due to the experimentally observed temporal smearing of dopamine signals [55, 56]. In real neural circuits, however, such imprecision may arise at multiple processing stages.

Firing rates of 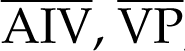, and VTA are described by the following equations:

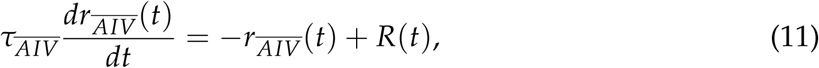

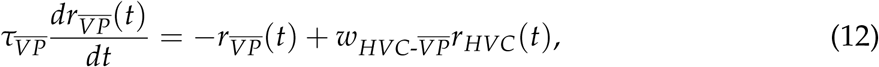

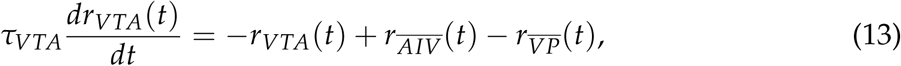

where the 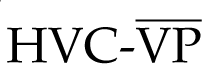 synaptic weight 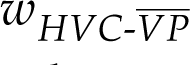 is a learnable parameter representing the reward prediction that is initialized to the true expected reward before learning for each task. 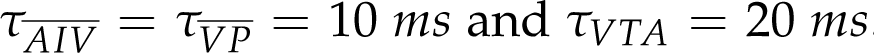. These firing rates and the 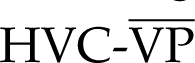 synaptic weights are defined relative to a baseline, and thus can be either positive or negative.

The values of reward are defined practically for each task: In Figure 2, we assume the baseline pitch is lower than the target pitch and leads to a reward of −1 (we define the reward as non-positive to be conceptually consistent with the fact that bird song is compared with an optimal template). When only the pitch-increasing LMAN neuron fires, improving the song performance, the reward is 0. When only the pitch-decreasing LMAN neuron fires, impairing the song performance, the reward is −2. In Figures 4 and 6, we assume the baseline reward is −1 if the LMAN neuron does not fire. If the LMAN neuron fires at a note that improves the song performance, the reward is 0. If the LMAN neuron fires at a note that impairs the song performance, the reward is −2. Since the reward prediction is subtracted from the reward, an arbitrary shift of the values of the reward has no effect on song performance. The effect of LMAN spikes should be considered as increasing or decreasing the reward from the baseline. For example, in Figure 3C, we assume the spike of the first LMAN neuron increases the reward by 1 and the spike of the second LMAN neuron increases the reward by 3 (chosen to be asymmetric for illustration purposes). In the auditory feedback task (Figure 5), the value of reward is binary and equals −1 if the pitch within the target syllable crosses the threshold for triggering noisy feedback, and 0 if the pitch does not cross the threshold. In Figure 7, the value of reward is proportional to the pitch and normalized between −2 and 0, with the baseline reward set to −1 when none of the LMAN neurons fire.

### Neural Circuit Model: Learning Rule

During learning, the 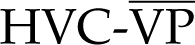 synapses receive dopaminergic input from the VTA neurons and undergo plasticity to induce critic learning (i.e., updating of reward prediction) according to the following critic learning rule:

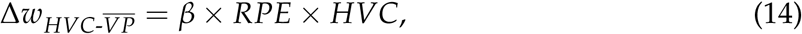

where *β* is a constant update speed. The HVC-MSN synapse undergoes plasticity to induce actor learning (i.e., a change in the LMAN spiking probability) according to the following actor learning rule:

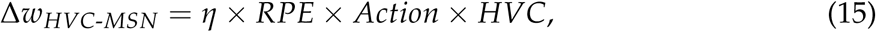

where *η* is a constant learning rate.

The *RPE* term corresponding to each note is represented by the VTA firing rate at the end of the note, multiplied by a scaling factor of 10 (e.g., if the reward is 1 and the reward prediction is 0, the VTA firing rate at the end of the note is approximately 0.1).

### Simulations

Within each learning trial, the activity in the actor, the activity in the critic, and the synaptic weight updates are simulated sequentially: 1) actions in the actor component (LMAN spiking activity) determine the value of reward for the critic component; 2) the critic component generates the dopamine signal (VTA activity); 3) plasticity rules adjust HVC-MSN weights in the actor and HVC-VP weights in the critic. The dynamical equations describing the neural circuit model were numerically solved using the Euler method with a time step of 0.05 *ms*.

### Simplified Model of LMAN firing probability

All simulations in the paper are performed in the actual neural circuit model with one exception: in Figure 7D-F, to control the maximum firing probability of the LMAN neurons, *p_max_*, we used a simplified actor with action probability *p* that mimicked the behavior of the neural circuit model (Figure 2B). Specifically, for these panels, we directly modeled the LMAN spiking probability as being the following function of the HVC-MSN weight *w*:

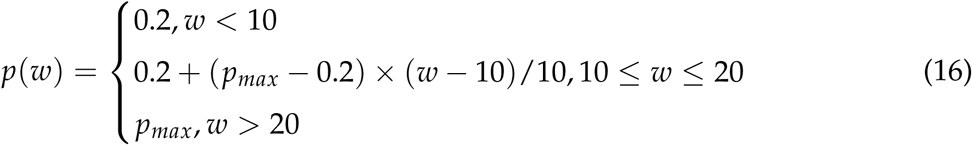

### Analysis of Experimental Data

We obtained an update speed of about *β* = 0.3 based on the data reported in [53]. They reported the coefficients of a linear regression fit of the reward history to the dopamine signal (see Fig. 4d in their paper). According to the critic learning rule (Eq. 4), the reward prediction at trial *t* can be shown to be (Supplemental Information, Eq. S80):

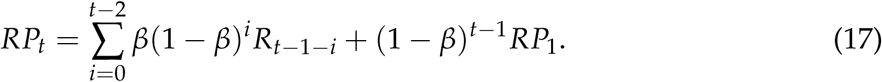

Thus, the reward prediction error (represented by the dopamine signal) at trial *t* is related to the reward history according to the following equation:

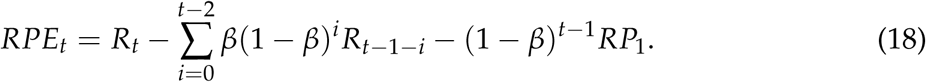

The coefficients corresponding to the current trial *t* and previous trials *t* − 1, *t* − 2, *t* − 3, etc., are 1, −*β*, −*β*(1 − *β*), −*β*(1 − *β*)^2^, etc. We fit the reported coefficients to the model to obtain a best fit *β* value of approximately 0.3.

We obtained the song learning trajectory in the CAF experiment (Figure 5B) from [38]. They reported an example trajectory of the average pitch of the target syllable during one day’s learning (see Fig. 1C in their paper). The original, raw trial-by-trial data were lost. We reconstructed the data based on the digital version of the data figure, in which trials close in time are grouped together into bins due to the limited resolution of the digitized format.

## Acknowledgments

We would like to express our sincere gratitude to Alex Sood, Alireza Alemi, Avinash Avinash, Ben Lankow, Jay Bhasin, Jiacheng Xu, and Lindsey Brown for their invaluable discussions and insights; Aaron Andalman for sharing his data; and Hannah Eum and Margarita O’Leary for their support. This work is funded by the Simons Collaboration on the Global Brain (M.G., M.F.).

## Author Contributions

Y.W., M.F., and M.G. designed the study. Y.W. performed the modeling and data analysis. Y.W., J.K., M.F., and M.G. wrote the manuscript.

## Code and Data Availability

Model code and experimental data are available on GitHub: https://github.com/goldman-lab/Bird-song-actor-critic-circuit-model.

