## Supplemental Information for "Constraints for spatially and temporally precise learning in a neural circuit model of reinforcement learning"

#### 1 Mathematical analysis of the RPE learning rule

In this section, we first show that the RPE learning rule implements stochastic gradient ascent in a major class of reinforcement learning problems known as finite Markov Decision Processes (MDPs). We show that the gradient represents the difference in value of choosing one action versus the other actions. We then relate this theoretical result to the neural circuit model of song learning.

**RPE learning rule implements stochastic gradient ascent**

In a finite Markov Decision Process, at each time step $t$, a learning agent receives information about the state of the environment, $S_t \in \mathcal{S}$, where $\mathcal{S}$ is a finite set of states. The agent selects an action $A_t$ from a finite set of actions $\mathcal{A}$. The action leads to a numerical reward $R_t \in \mathcal{R} \subset \mathbb{R}$ where $\mathcal{R}$ is a finite set of rewards, and the environment transitions into a new state $S_{t+1}$ at the next time point. We restrict ourselves to episodic problems in which each episode starts at time $t = 0$ from an initial state $S_0$ drawn from a fixed initial-state distribution $\rho_0$ and ends at a finite time point $T$. In the following, we derive that, for the episodic finite MDP, the RPE learning rule implements stochastic gradient ascent.

For any state $S_t$ within an episode extending from time 0 to time $T$, the sum of the future rewards starting from state $S_t$ is called the return, denoted by $G(S_t)$:

$$G(S_t) := \sum_{i=t}^{T} R_i. \tag{S1}$$

The expected return when starting from state $S_t$ is called the state value, denoted by $v(S_t)$:

$$v(S_t) := \mathbb{E}[G(S_t)]. \tag{S2}$$

The goal of learning is to maximize the expected return, averaged over all possible initial states $S_0$, which we refer to as the target function $J$:

The sum of the future rewards after choosing an action $a$ at state $S_t$ is denoted by $G(S_t, a)$:

$$G(S_t, a) := \sum_{i=t}^{T} R_i \text{ , } \textit{if } A_t = a. \tag{S3}$$

The expected return after choosing an action $a$ at state $S_t$ is called the state-action value, denoted by $q(S_t, a)$:

$$q(S_t, a) := \mathbb{E}[G(S_t, a)]. \tag{S4}$$

At each state $S_t$, the learning agent chooses the action according to a probability distribution, called the policy of the agent. Specifically, we use $p(a|S_t)$ to denote the probability of choosing action $a$ at state $S_t$. The state value and the state-action value are related by:

$$v(S_t) = \sum_{a} q(S_t, a) p(a|S_t). \tag{S5}$$

A family of reinforcement learning algorithms called policy gradient methods assumes that the action probabilities $p(a|S_t)$ are determined by the action preferences $w(a|S_t)$ according to a softmax distribution:

$$p(a|S_t, \boldsymbol{w}) = \frac{e^{w(a|S_t)}}{\sum_{b \in \mathcal{A}} e^{w(b|S_t)}}, \tag{S6}$$

where the vector $\boldsymbol{w}$ denotes all the action preferences. The target function $J$ can can then be written in the following form:

$$J(\boldsymbol{w}) := \mathbb{E}_{S_0 \sim \rho_0, p_{\boldsymbol{w}}}[G(S_0)], \tag{S7}$$

where $p_{\boldsymbol{w}} := p(\cdot|S_t, \boldsymbol{w})$ denotes the policy. In policy gradient methods, the learning goal is achieved by updating the action preferences $\boldsymbol{w}$ along the gradient of the target function. According to the policy gradient theorem [1, 2], the gradient can be written as:

$$\nabla_{\boldsymbol{w}} J(\boldsymbol{w}) = \mathbb{E}_{S_0 \sim \rho_0, p_{\boldsymbol{w}}} \left[ \sum_{t=0}^{T} \boldsymbol{g}(S_t) \right],$$

where $\boldsymbol{g}(S_t)$ denotes the contribution to the policy gradient from a visit to state $S_t$:

$$\boldsymbol{g}(S_t) = \sum_{a} q(S_t, a) \nabla_{\boldsymbol{w}} p(a|S_t, \boldsymbol{w}). \tag{S8}$$

By adding terms that evaluate to 0, we can further obtain that:

$$\boldsymbol{g}(S_t) = \sum_a \big[q(S_t,a) - v(S_t)\big] \nabla_{\boldsymbol{w}} p(a|S_t,\boldsymbol{w}) \tag{S9}$$

$$= \sum_a \big[q(S_t,a) - v(S_t)\big] \nabla_{\boldsymbol{w}} p(a|S_t,\boldsymbol{w}) + \boldsymbol{K}\big[v(S_t) - v(S_t)\big]. \tag{S10}$$

Above, the subtraction of $v(S_t)$ from $q(S_t, a)$ does not change the equality because

$$\sum_a v(S_t) \nabla_{\boldsymbol{w}} p(a|S_t,\boldsymbol{w}) = v(S_t) \nabla_{\boldsymbol{w}} \sum_a p(a|S_t,\boldsymbol{w}) = 0, \tag{S11}$$

and we add a term $\boldsymbol{K}\big[v(S_t) - v(S_t)\big] \equiv 0$, where each element of the vector $\boldsymbol{K}$ (corresponding to each action preference $w(a)$) can be any arbitrary constant.

Now we write $v(S_t)$ in two equivalent forms:

$$v(S_t) = \sum_a p(a|S_t,\boldsymbol{w}) q(S_t,a) = \sum_a p(a|S_t,\boldsymbol{w}) v(S_t). \tag{S12}$$

Thus we have:

$$\boldsymbol{g}(S_t) = \sum_a \big[q(S_t,a) - v(S_t)\big] \nabla_{\boldsymbol{w}} p(a|S_t,\boldsymbol{w}) + \boldsymbol{K}\big[v(S_t) - v(S_t)\big] \tag{S13}$$

$$= \sum_a p(a|S_t,\boldsymbol{w}) \left[q(S_t,a) - v(S_t)\right] \frac{\nabla_{\boldsymbol{w}} p(a|S_t,\boldsymbol{w})}{p(a|S_t,\boldsymbol{w})} \tag{S14}$$

$$+ \boldsymbol{K} \left[\sum_a p(a|S_t,\boldsymbol{w}) q(S_t,a) - \sum_a p(a|S_t,\boldsymbol{w}) v(S_t)\right] \tag{S15}$$

$$= \sum_a p(a|S_t,\boldsymbol{w}) \left[q(S_t,a) - v(S_t)\right] \left[\frac{\nabla_{\boldsymbol{w}} p(a|S_t,\boldsymbol{w})}{p(a|S_t,\boldsymbol{w})} + \boldsymbol{K}\right]. \tag{S16}$$

This can be written as the expectation under the policy $p(\cdot|S_t, \boldsymbol{w})$:

$$\boldsymbol{g}(S_t) = \mathbb{E}_{A_t \sim p(\cdot|S_t,\boldsymbol{w})} \Big[ \left[q(S_t,A_t) - v(S_t)\right] \Big[\frac{\nabla_{\boldsymbol{w}} p(A_t|S_t,\boldsymbol{w})}{p(A_t|S_t,\boldsymbol{w})} + \boldsymbol{K}\Big]\Big], \tag{S17}$$

as long as the elements of the vector $\boldsymbol{K}$ are constants independent of the sampled action $A_t$.

In a given episode, the return is $G(S_t, A_t)$ when action $A_t$ is chosen at state $S_t$. Replacing $q(S_t, A_t)$ by $G(S_t, A_t)$, we get an unbiased stochastic sample of the per-visit policy-gradient contribution $\boldsymbol{g}(S_t)$:

$$\left[G(S_t,A_t) - v(S_t)\right] \left[\frac{\nabla_{\boldsymbol{w}} p(A_t|S_t,\boldsymbol{w})}{p(A_t|S_t,\boldsymbol{w})} + \boldsymbol{K}\right]. \tag{S18}$$

For any action $a$, from Eq. S6, we have that:

$$\frac{\nabla_{w(a|S_t)} p(A_t|S_t, \boldsymbol{w})}{p(A_t|S_t, \boldsymbol{w})} = \mathbb{1}_{A_t=a} - p(a|S_t), \tag{S19}$$

where $\mathbb{1}_{A_t=a} = 1$ if the chosen action $A_t$ is $a$ and $\mathbb{1}_{A_t=a} = 0$ otherwise. By choosing $K(a) = p(a|S_t)$ for each action $a$, we obtain the RPE learning rule:

$$\Delta w(a|S_t) \propto \left[G(S_t, A_t) - v(S_t)\right] \times \mathbb{1}_{A_t=a}. \tag{S20}$$

In the above, $G(S_t, A_t)$ is the actual sum of the future rewards received by the learning agent after choosing action $A_t$ at state $S_t$, and $v(S_t)$ is the expectation value of the sum of the future rewards at state $S_t$. Thus, $G(S_t, A_t) - v(S_t)$ is the reward prediction error (RPE).

In practice, the true state value, $v(S_t)$ typically needs to be learned and approximated. In this case, the action preferences are updated according to the following equation:

$$\Delta w(a|S_t) \propto \left[G(S_t, A_t) - V(S_t)\right] \times \mathbb{1}_{A_t=a}, \tag{S21}$$

where $V(S_t)$ is an estimate of the state value $v(S_t)$. When $V(S_t) = v(S_t)$, this update is an unbiased stochastic estimator of the policy gradient. When $V(S_t) \neq v(S_t)$, the update contains an additional bias term and therefore does not necessarily follow the true policy gradient, which can cause the interference problems shown in the main text.

In the derivation above, the softmax distribution can include a temperature parameter $h$:

$$p(a|S_t, \boldsymbol{w}) = \frac{e^{hw(a|S_t)}}{\sum_{b\in\mathcal{A}} e^{hw(b|S_t)}}. \tag{S22}$$

However, it can be easily shown that the temperature parameter only appears as a constant scaling factor in the gradient and thus can be merged with the learning rate. As a result, the derivation above assumes a temperature parameter of 1 for simplicity but applies generally. In conclusion, the derivation above demonstrates that, when the reward prediction is accurate, the RPE learning rule implements stochastic gradient ascent in episodic finite Markov Decision Processes.

**Application to the multi-armed bandit problem.**

In the main text, we consider the problem of LMAN neurons learning to spike or not spike at each note to be an instantiation of the multi-armed bandit problem. For a multi-armed bandit problem, each trial can be treated as a one-step episode with only one state. Thus, the return $G$ reduces to the immediate reward $R$, and the state value $v$ is simply the expectation value of the reward $\mathbb{E}[R]$. In this case, the RPE learning rule can be written as:

$$\Delta w(a) \propto (R - \mathbb{E}[R]) \times \mathbb{1}_{A=a}. \tag{S23}$$

As noted above, typically the expectation value of the reward $\mathbb{E}[R]$ needs to be learned. In this case, the action preferences are updated according to:

$$\Delta w(a) \propto (R - RP) \times \mathbb{1}_{A=a}, \tag{S24}$$

where $RP$ denotes the learned estimate of the reward prediction. When the reward prediction deviates from the true expectation value of the reward, which can be caused by the interference effects shown in the main text, incorrect learning can occur.

**RPE learning rule approximately performs stochastic gradient ascent in the neural circuit model of song learning**

In the neural circuit model of song learning, the LMAN neuron makes a binary decision of whether to spike or not spike. Using $Action = 1$ and $Action = 0$ to denote whether the LMAN neuron spikes or not, the RPE learning rule can be written as:

$$\Delta w \propto (R - RP) \times Action. \tag{S25}$$

The LMAN spiking probability, $p(Spike)$, is determined by the HVC-MSN synaptic weight, $w$. If the relationship between $p(Spike)$ and $w$ follows a logistic function:

$$p(Spike|w) = \frac{1}{1 + e^{-k(w-w_0)}}, \tag{S26}$$

where $k$ is a constant temperature parameter (that can be assumed to be 1 without loss of generality), then the RPE learning rule implements stochastic gradient ascent, following the same derivation process shown above, since

$$\frac{\nabla_w p(A|w)}{p(A|w)} = \mathbb{1}_{A=Spike} - p(Spike|w). \tag{S27}$$

The actual relationship between the LMAN firing probability and the HVC-MSN synaptic weight in the neural circuit model does not strictly follow a logistic function, but only has an approximately sigmoidal shape (Figure 2B). Nevertheless, empirical simulations show that the RPE learning rule does approximately perform stochastic gradient ascent (Figure 3C).

**RPE learning rule strengthens actions with higher expected reward**

As the derivation above shows, the RPE learning rule implements stochastic gradient ascent (assuming the reward prediction is accurate). This means that, conditional on visiting state $S_t$, the expected change of the action preference $\Delta w(a|S_t)$ is proportional to the corresponding component of the per-visit policy-gradient contribution $\boldsymbol{g}(S_t)$. Here, we show that this represents the value difference between choosing the action $a$ and not choosing $a$.

We define the value of not choosing action $a$ at state $S_t$ as:

$$q(S_t, \bar{a}) := \mathbb{E}[G(S_t, A_t \neq a)] = \frac{\sum_{b \neq a} p(b|S_t) q(S_t, b)}{\sum_{b \neq a} p(b|S_t)} = \frac{\sum_{b \neq a} p(b|S_t) q(S_t, b)}{1 - p(a|S_t)}, \tag{S28}$$

where, for simplicity, we have omitted writing the parameter $\boldsymbol{w}$ in $p(a|S_t, \boldsymbol{w})$. Thus,

$$\mathbb{E}[\Delta w(a|S_t)] \propto \mathbb{E}\big[[G(S_t, A_t) - v(S_t)] \times \mathbb{1}_{A_t=a}\big] \tag{S29}$$

$$= p(a|S_t)[q(S_t, a) - v(S_t)] \tag{S30}$$

$$= p(a|S_t)\big[q(S_t, a) - \sum_b p(b|S_t) q(S_t, b)\big] \tag{S31}$$

$$= p(a|S_t)\big[q(S_t, a) - p(a|S_t) q(S_t, a) - \sum_{b \neq a} p(b|S_t) q(S_t, b)\big] \tag{S32}$$

$$= p(a|S_t)[1 - p(a|S_t)]\Big[q(S_t, a) - \frac{\sum_{b \neq a} p(b|S_t) q(S_t, b)}{1 - p(a|S_t)}\Big] \tag{S33}$$

$$= p(a|S_t)[1 - p(a|S_t)][q(S_t, a) - q(S_t, \bar{a})]. \tag{S34}$$

From the final term in the above equation, the RPE learning rule will strengthen the preference of an action in proportion to the difference between the value of taking this action and the value of not taking this action.

We note that this result does not require that the action probabilities $p(a|S_t)$ are determined by action preferences $w(a|S_t)$ according to the softmax distribution (Eq. S6). It therefore provides a more general interpretation of the RPE learning rule. This interpretation is relevant to models such as the neural circuit model of song learning, in which the relationship between the LMAN spiking probability and the HVC-MSN synaptic weight deviates from the exact logistic form (Figure 2B) and the learning rule therefore only approximates stochastic gradient ascent.

### 2 Temporal interference causes over-prediction and unlearning

In the main text, we showed that temporal imprecision of the RPE signaling can cause interference and incorrect learning. In this section, we show that it can also cause the reward prediction to exceed the actual expected reward, which we call the over-prediction problem. Furthermore, when the update speed of reward prediction is very slow, the over-prediction problem causes significant unlearning, in which the LMAN spiking probability drops after it learns to increase. The over-prediction and unlearning problems can be mitigated by a relatively fast update speed of reward prediction. To illustrate this, we consider the same learning scenario as in Figure 6 of the main text, in which one LMAN neuron learns at five consecutive notes. In the main text, we showed the incorrect learning problem at note 5. Here, we focus on issues in learning the first four notes.

We first demonstrate the over-prediction problem (Figure S1A-C). We assume the reward predictions are updated according to:

$$RP_i(t+1) = RP_i(t) + \beta RPE_i(t), \tag{S35}$$

where $i = 1, \ldots, 4$ indicates the four notes, and $t$ indicates the number of learning trials ($t$ starts from 1). As one example, we initialize the reward predictions at trial $t = 1$ at the four consecutive notes to be $RP_i(1) = -1$, and constantly provide a reward of 0 for all notes.

If the RPE signal is temporally precise,

$$RPE_i(t) = R_i(t) - RP_i(t) = 0 - RP_i(t), \tag{S36}$$

then all four reward predictions will follow the same geometric decay towards the true expected reward of 0 (Figure S1A):

$$RP_i(t) = -(1-\beta)^{t-1}, \tag{S37}$$

for all $i$. When $\beta$ is small, this geometric decay can be approximated by an exponential decay:

$$RP_i(t) \approx -e^{-\beta(t-1)}. \tag{S38}$$

If instead the RPE signal is temporally imprecise such that the RPE signal at one note extends across the future notes (Figure S1B), we calculate the RPE at notes $i > 1$ by

$$RPE_i(t) = R_i(t) - RP_i(t) + \gamma RPE_{i-1}(t) = 0 - RP_i(t) + \gamma RPE_{i-1}(t), \tag{S39}$$

where $\gamma$ is a decay factor.

For the first note, which is not affected by RPE signals from the previous notes, we have:

$$RP_1(t) = -(1-\beta)^{t-1}. \tag{S40}$$

To simplify notation, we denote

$$b = 1 - \beta. \tag{S41}$$

Thus,

$$RP_1(t) = -b^{t-1}, \quad RPE_1(t) = b^{t-1}. \tag{S42}$$

For the second note, the reward prediction is updated as:

$$RP_2(t+1) = RP_2(t) + \beta RPE_2(t) \tag{S43}$$

$$= RP_2(t) + \beta(0 - RP_2(t) + \gamma RPE_1(t)) \tag{S44}$$

$$= bRP_2(t) + \beta\gamma RPE_1(t) \tag{S45}$$

$$= bRP_2(t) + \beta\gamma b^{t-1}. \tag{S46}$$

Solving this discrete equation gives:

$$RP_2(t) = b^{t-1}\left(-1 + \frac{\beta\gamma}{b}(t-1)\right). \tag{S47}$$

Substituting $b = 1 - \beta$ back into the equation gives:

$$RP_2(t) = (1-\beta)^{t-1}\left(-1 + \frac{\beta\gamma}{1-\beta}(t-1)\right). \tag{S48}$$

Therefore,

$$RPE_2(t) = -RP_2(t) + \gamma RPE_1(t) \tag{S49}$$

$$= (1+\gamma)b^{t-1} - \beta\gamma(t-1)b^{t-2}. \tag{S50}$$

Similarly, for the third note:

$$RP_3(t+1) = bRP_3(t) + \beta\gamma RPE_2(t) \tag{S51}$$

$$= bRP_3(t) + \beta\gamma(1+\gamma)b^{t-1} - (\beta\gamma)^2(t-1)b^{t-2}. \tag{S52}$$

Solving this discrete equation gives:

$$RP_3(t) = b^{t-1}\Big[-1 + \frac{\beta\gamma}{b}(1+\gamma)(t-1) - \left(\frac{\beta\gamma}{b}\right)^2 \frac{(t-1)(t-2)}{2}\Big]. \tag{S53}$$

Substituting $b = 1-\beta$ back into the equation gives:

$$RP_3(t) = (1-\beta)^{t-1}\Big[-1 + \frac{\beta\gamma}{1-\beta}(1+\gamma)(t-1) - \left(\frac{\beta\gamma}{1-\beta}\right)^2 \frac{(t-1)(t-2)}{2}\Big]. \tag{S54}$$

Therefore,

$$RPE_3(t) = -RP_3(t) + \gamma RPE_2(t) \tag{S55}$$

$$= (1+\gamma+\gamma^2)b^{t-1} - \beta\gamma(1+2\gamma)(t-1)b^{t-2} + (\beta\gamma)^2\frac{(t-1)(t-2)}{2}b^{t-3}. \tag{S56}$$

For the fourth note:

$$RP_4(t+1) = bRP_4(t) + \beta\gamma RPE_3(t) \tag{S57}$$

$$= bRP_4(t) + \beta\gamma(1+\gamma+\gamma^2)b^{t-1} \tag{S58}$$

$$- (\beta\gamma)^2(1+2\gamma)(t-1)b^{t-2} + (\beta\gamma)^3\frac{(t-1)(t-2)}{2}b^{t-3}. \tag{S59}$$

Solving this discrete equation gives:

$$RP_4(t) = b^{t-1}\Big[-1 + \frac{\beta\gamma}{b}(1+\gamma+\gamma^2)(t-1) - \left(\frac{\beta\gamma}{b}\right)^2(1+2\gamma)\frac{(t-1)(t-2)}{2} \tag{S60}$$

$$+ \left(\frac{\beta\gamma}{b}\right)^3 \frac{(t-1)(t-2)(t-3)}{6}\Big]. \tag{S61}$$

Substituting $b = 1-\beta$ back into the equation gives:

$$RP_4(t) = (1-\beta)^{t-1}\Big[-1 + \frac{\beta\gamma}{1-\beta}(1+\gamma+\gamma^2)(t-1) \tag{S62}$$

$$- \left(\frac{\beta\gamma}{1-\beta}\right)^2(1+2\gamma)\frac{(t-1)(t-2)}{2} + \left(\frac{\beta\gamma}{1-\beta}\right)^3 \frac{(t-1)(t-2)(t-3)}{6}\Big]. \tag{S63}$$

In summary,

$$RP_1(t) = -(1-\beta)^{t-1}. \tag{S64}$$

$$RP_2(t) = (1-\beta)^{t-1}\left(-1 + \frac{\beta\gamma}{1-\beta}(t-1)\right). \tag{S65}$$

$$RP_3(t) = (1-\beta)^{t-1}\Big[-1 + \frac{\beta\gamma}{1-\beta}(1+\gamma)(t-1) - \left(\frac{\beta\gamma}{1-\beta}\right)^2 \frac{(t-1)(t-2)}{2}\Big]. \tag{S66}$$

$$RP_4(t) = (1-\beta)^{t-1}\Big[-1 + \frac{\beta\gamma}{1-\beta}(1+\gamma+\gamma^2)(t-1) \tag{S67}$$

$$- \left(\frac{\beta\gamma}{1-\beta}\right)^2 (1+2\gamma)\frac{(t-1)(t-2)}{2} + \left(\frac{\beta\gamma}{1-\beta}\right)^3 \frac{(t-1)(t-2)(t-3)}{6}\Big]. \tag{S68}$$

These solutions show that temporal imprecision introduces additional terms, of the form of a geometric decay times a polynomial in $t$, on top of the original geometric decay. The additional terms can push the reward prediction above the true expected reward of 0, thereby producing over-prediction. Figure S1B shows an example in which $\gamma = 1$. Figure S1C demonstrates the over-prediction problem by simulating the critic component of the neural circuit model of song learning (Methods) for the case of fixed rewards. .

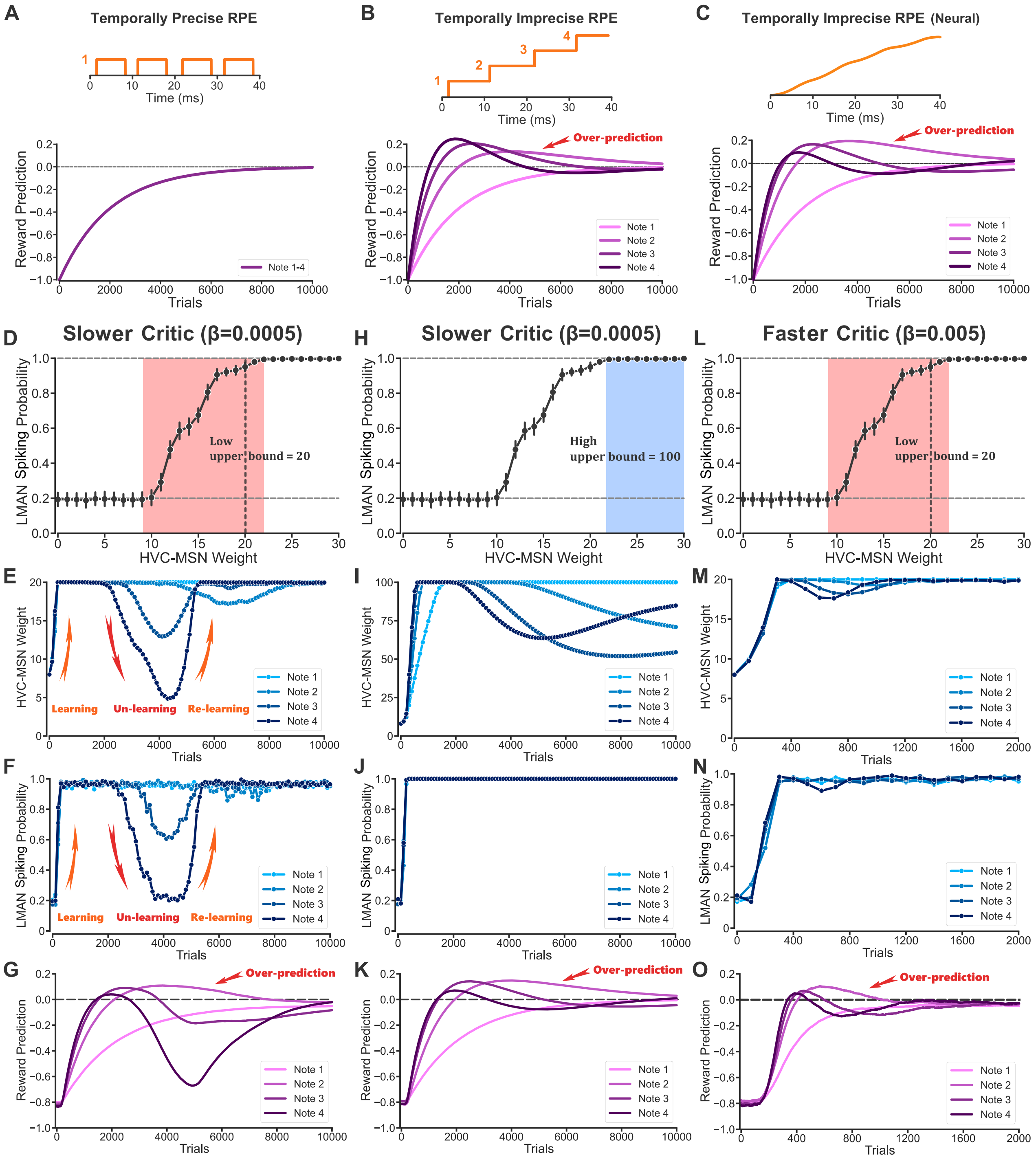


**Figure S1: Temporal interference causes over-prediction and unlearning. (A)** Reward predictions follow a geometric decay to the true reward value of zero if the RPE signal is temporally precise. **(B)** Temporal imprecision causes over-prediction, illustrated for the case of fixed rewards and no decay of the RPE signals. **(C)** As in B, but using the equations corresponding to the song-learning critic (Eqs. 11-13, Methods) for the updating of reward prediction and calculation of reward prediction error. **(D-G)** Temporal interference causes over-prediction and unlearning in the full neural circuit model of song learning. (D) LMAN spiking probability as a function of the HVC-MSN weight. A hard upper bound of 20 is imposed on the weight in this example. This upper bound is close to the boundary of the range over which changes in the HVC-MSN weight change the LMAN spiking probability (i.e., the 'dynamic range', red background). (E) HVC-MSN weight across learning. (F) LMAN spiking probability across learning. (G) Reward prediction across learning. **(H-K)** The unlearning problem can be mitigated by letting the upper bound on the HVC-MSN weight be much higher than the dynamic range. Same as panels D-G but with a high upper bound (100) for the synaptic weight. The blue background represents the non-dynamic range of the HVC-MSN weight that contains this bound. **(L-O)** The unlearning problem can be mitigated by using a relatively fast critic ($\beta = 0.005$ in this example versus $\beta = 0.0005$ in the previous two examples). All data points show the average of 100 repetitions. Error bars in panels D, H, and L indicate standard deviation across 100 repetitions (not shown in the other panels for visual clarity).

Next, we simulate the full song-learning network of Figure 6 of the main text and show that when the update speed of reward prediction is very slow, the over-prediction of the reward can cause significant unlearning, in which the LMAN spiking probability drops after it learns to increase (Figure S1D-G). In this example, the update speed $\beta$ is set to be 0.0005, and we impose a strict upper bound of 20 on the HVC-MSN weight, close to the boundary of the range over which changes in the HVC-MSN weight change the LMAN spiking probability, which we refer to as the 'dynamic range' (Figure S1D, red background). The steps underlying this unlearning are as follows: Over-prediction means the predicted reward is higher than the true expectation value of the reward, which leads to a negative RPE. This negative RPE causes the HVC-MSN weight to decrease (Figure S1E), which causes the LMAN spiking probability to decrease (Figure S1F). Since we assumed that spiking of this LMAN neuron increases reward, decreased probability of LMAN spiking leads to a decreased expectation value of the reward, causing an even more negative RPE and a further decrease of the HVC-MSN synaptic weight and expected reward. This positive feedback causes the LMAN spiking probability to drop substantially after it learns to increase. Due to the temporal imprecision of the RPE signaling, the negative RPE signals in earlier notes also smear to later notes (similar to the toy example of Figure S1C, in which the RPE signals were positive). This causes the unlearning problem to be more significant for the later notes

(Figure S1E-G).

The unlearning problem can be avoided if the upper bound of the HVC-MSN weight is much higher than the dynamic range (Figure S1H-K). In this case, the negative RPE caused by over-prediction causes a decrease of the weight (Figure S1I) but not the LMAN spiking probability (Figure S1J) since the weight remains in the non-dynamic range (Figure S1H, blue background). Alternatively, the unlearning problem can be mitigated if the update speed of reward prediction is relatively fast (Figure S1L-O) since the reward prediction approaches the actual expected reward quickly (note the time scale in panels M-O versus previous panels) and thus exerts little effect on the HVC-MSN weight.

### 3 Effect of the actor learning rate

A slower actor learning rate places a less stringent requirement on the speed of critic learning. In Figure 7 of the main text, the actor learning rate was chosen to achieve learning in the model on the 1,000-trial timescale of a day of natural birdsong learning [3]. However, we can use a slower actor learning rate and a larger number of learning trials to achieve approximately the same amount of incorrect learning with a slower critic update speed (Figure S2).

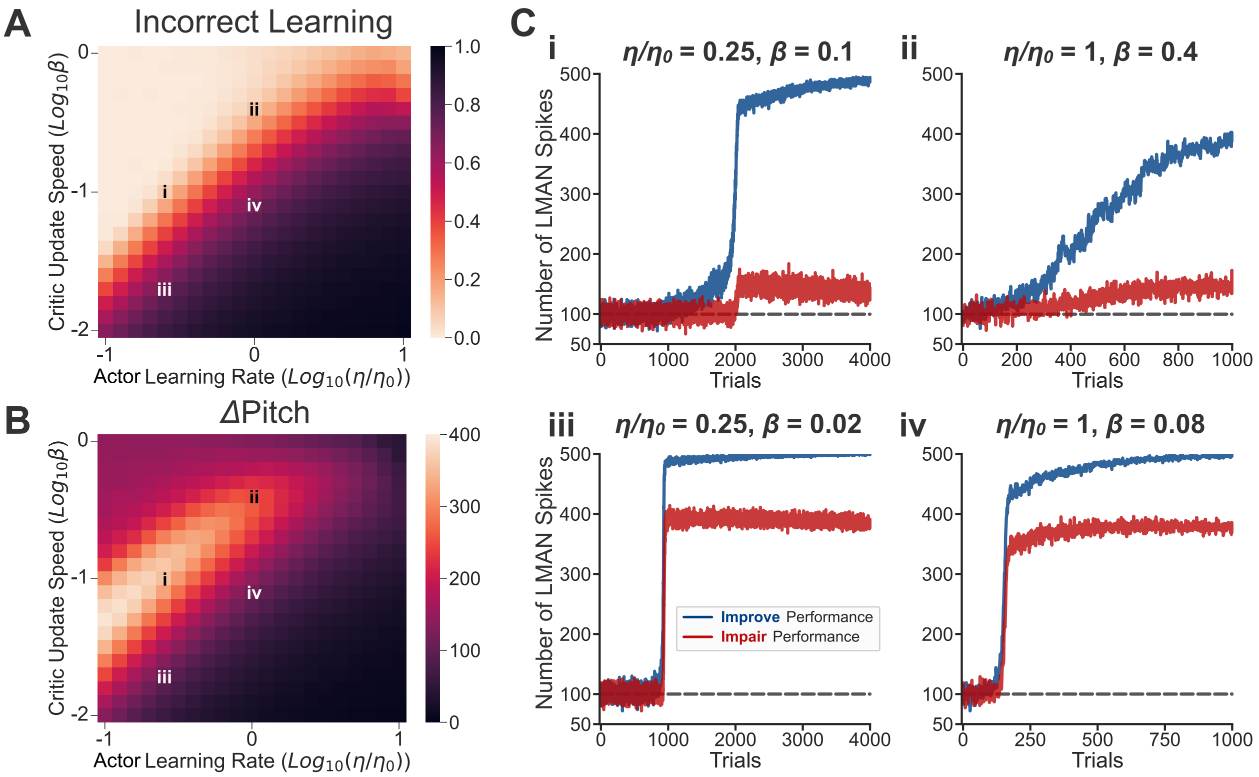


**Figure S2: A slower actor learning rate places a less stringent requirement on the speed of critic learning. (A-C)** Effect of the reward prediction update speed $\beta$ and the actor learning rate $\eta$ on the extent of incorrect learning (A), the overall learned pitch increase (B), and the change in the number of LMAN spikes (C), respectively. Subpanels (i-iv) in panel C correspond to different parameter settings in panels A and B. $\eta_0 = 15$ represents the actor learning rate used in Figure 7 of the main text.

### 4 Effect of the number of LMAN neurons

In the main text, we showed that interference across motor channels can lead to an excessively positive RPE and cause incorrect learning when the update speed of reward prediction is slow (Figure 7). Here, we show that the extent of this problem critically depends on the number of LMAN neurons involved in learning.

According to the RPE learning rule:

$$\Delta w = \eta(R - RP) \times Action. \tag{S69}$$

When the reward prediction $RP$ is accurate:

$$RP = \mathbb{E}[R] = p\mathbb{E}[R|Action = 1] + (1-p)\mathbb{E}[R|Action = 0], \tag{S70}$$

where $p$ is the LMAN spiking probability. The expected change of the HVC-MSN synaptic weight across trials for each LMAN neuron is

$$\begin{aligned} \mathbb{E}[\Delta w] &= \eta p \Big[\mathbb{E}[R|Action = 1] - p\mathbb{E}[R|Action = 1] - (1-p)\mathbb{E}[R|Action = 0]\Big] && \text{(S71)} \\ &= \eta p(1-p)\Big[\mathbb{E}[R|Action = 1] - \mathbb{E}[R|Action = 0]\Big]. && \text{(S72)} \end{aligned}$$

We can see that the expected change of synaptic weight is proportional to the term $\mathbb{E}[R|Action = 1] - \mathbb{E}[R|Action = 0]$, which is the value difference between LMAN spiking ($Action = 1$) and not spiking ($Action = 0$). This can be considered as the 'driving force of learning' for each LMAN neuron, which depends on the effect that the spike of this LMAN neuron has on the reward received. We consider the case in which the pitch starts below its target value, so that it needs to be increased. If we assume the spike of each LMAN neuron changes the reward by 1 unit, the driving force of learning for each LMAN neuron is $+1$ (if it increases the pitch and improves the song performance) or $-1$ (if it decreases the pitch and impairs the song performance).

Now consider what happens when the reward prediction is inaccurate. Specifically, when the expected reward increases but the reward prediction is slow to catch up, there will be a bias in the reward prediction:

$$RP = \mathbb{E}[R] - Bias, \tag{S73}$$

where the *Bias* term is greater than 0. Since the bias comes from the overall change of behavioral performance contributed by all of the LMAN neurons, it can dominate learning when the update speed of reward prediction is slow and the number of LMAN neurons is large.

The RPE learning rule can now be written as:

$$\Delta w = \eta(R - \mathbb{E}[R] + Bias) \times Action \tag{S74}$$

$$= \eta(R - \mathbb{E}[R]) \times Action + \eta Bias \times Action. \tag{S75}$$

The expected change of synaptic weight across trials for each LMAN neuron becomes:

$$\mathbb{E}[\Delta w] = \eta p(1-p)\Big[\mathbb{E}[R|Action = 1] - \mathbb{E}[R|Action = 0]\Big] + \eta p Bias, \tag{S76}$$

which contains the same driving force of learning as above plus a positive bias term. The incorrect learning problem occurs for the performance-impairing neurons when such positive bias dominates over the driving force of learning for these neurons, as can especially occur when the number of LMAN neurons is large. Indeed, Figure S3 shows that the incorrect learning problem only occurs when there are a sufficient number of neurons, and an increasingly fast reward prediction update speed is required to avoid incorrect learning, and to achieve the maximum amount of learned pitch increase, with a larger number of neurons.

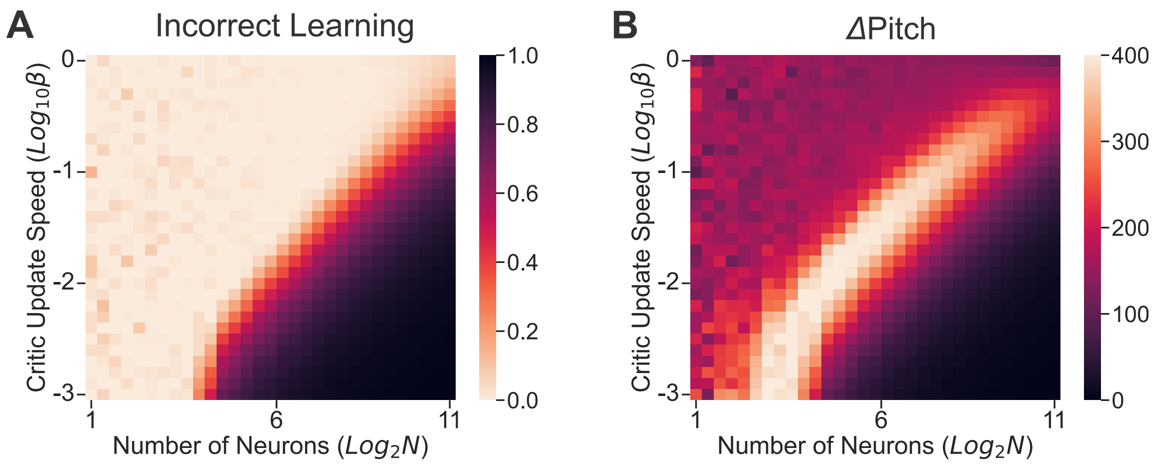


**Figure S3: Effect of the number of LMAN neurons.** **(A)** Effect of critic update speed $\beta$ and the number of LMAN neurons $N$ on the extent of incorrect learning. **(B)** Effect of critic update speed $\beta$ and the number of LMAN neurons $N$ on the overall learned pitch increase. Both panels show the average results across 100 repetitions, measured at trial $1,000$ in the same pitch learning scenario as Figure 7 of the main text, in which half of the LMAN neurons increase the pitch and improve performance while the other half decrease the pitch and impair performance. To compare across different numbers of neurons, we assume the maximum change of pitch ranges from $-500$ to 500 regardless of the actual number of LMAN neurons. For example, the spike of each LMAN neuron changes the pitch by 1 unit when there are $1,000$ LMAN neurons and the spike of each LMAN neuron changes the pitch by 10 units when there are 100 LMAN neurons. As a result, the fewer LMAN neurons there are, the more each neuron affects the behavior. To compensate for the resulting increased driving force of learning for each neuron (Eq. S72, term in brackets) and maintain a fixed expected change in weight $\mathbb{E}[\Delta w]$, the learning rate $\eta$ is adjusted proportionally to the number of LMAN neurons: $\eta = \eta_0 \times (N/1,000)$, where $\eta_0 = 15$ is the learning rate used in Figure 7 of the main text, in which there are $1,000$ LMAN neurons. The maximum LMAN spiking probability $p_{max}$ is fixed at 1.

### 5 Update speed of reward prediction

In this section, we analyze the effect of the reward prediction update speed $\beta$ on the reward prediction error when the reward prediction is updated by:

$$RP_{t+1} = RP_t + \beta(R_t - RP_t). \tag{S77}$$

Specifically, we ask what the optimal choice of $\beta$ is if one wants to minimize the reward prediction error (i.e., maximize the accuracy of reward prediction).

We use the expectation of the squared RPE as the criterion for the accuracy of reward prediction and show that it faces a bias-variance tradeoff. The idealized analysis below suggests a range of $\beta$ that is much smaller than the values found in the main text to optimize overall song learning performance.

**The variance**

We first consider a simple scenario in which the reward is drawn from a static Gaussian distribution with mean 0 and variance $\sigma^2$. In previous examples, we typically assumed a baseline reward of $-1$ to be consistent with song learning, in which the reward is always non-positive. Here, to obtain general theoretical insights, we assume the baseline reward is 0 for simplicity.

Suppose the reward prediction starts with an accurate prediction, i.e., $RP_1 = 0$, and is not updated at all, i.e., $\beta = 0$. Then, $RP_t = 0 = \mathbb{E}[R_t]$ for all $t$. The expectation value of the squared reward prediction error can be obtained as:

$$\mathbb{E}[(R_t - RP_t)^2] = Var[R_t - RP_t] + (\mathbb{E}[R_t - RP_t])^2 = Var[R_t] = \sigma^2. \tag{S78}$$

At the other extreme, if $\beta = 1$, then $RP_t = R_{t-1}$ for all $t \geq 2$, i.e., the current prediction equals the previous reward. In this case:

$$\mathbb{E}[(R_t - RP_t)^2] = \mathbb{E}[(R_t - R_{t-1})^2] = Var[R_t - R_{t-1}] + (\mathbb{E}[R_t - R_{t-1}])^2 = 2\sigma^2. \tag{S79}$$

Intuitively, the expectation value of the squared RPE will be a monotonically increasing function of the update speed $\beta$ (with a faster update speed, the reward prediction will be more affected by the noisiness of the reward and deviate from the true expected reward). Quantitatively, for $t \geq 2$:

$$RP_t = \sum_{i=0}^{t-2} \beta(1-\beta)^i R_{t-1-i} + (1-\beta)^{t-1} RP_1 = \sum_{i=0}^{t-2} \beta(1-\beta)^i R_{t-1-i}. \tag{S80}$$

Thus we have:

$$\mathbb{E}[(R_t - RP_t)^2] = Var[R_t - RP_t] + \mathbb{E}[R_t - RP_t]^2 \tag{S81}$$

$$= Var[R_t] + Var[\sum_{i=0}^{t-2} \beta(1-\beta)^i R_{t-1-i}] - 2Cov(R_t, \sum_{i=0}^{t-2} \beta(1-\beta)^i R_{t-1-i}) \tag{S82}$$

$$= Var[R_t] + Var[\sum_{i=0}^{t-2} \beta(1-\beta)^i R_{t-1-i}] \tag{S83}$$

$$= \sigma^2 + \sum_{i=0}^{t-2} \beta^2 (1-\beta)^{2i} \sigma^2 \tag{S84}$$

$$= \left[1 + \frac{\beta}{2-\beta}[1 - (1-\beta)^{2(t-1)}]\right]\sigma^2. \tag{S85}$$

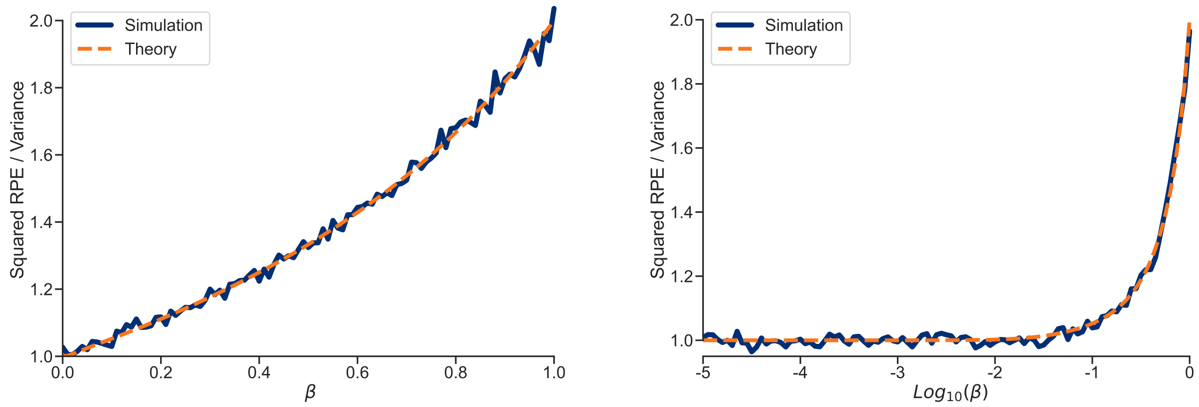


**Figure S4: Squared RPE (normalized by the reward variance $\sigma^2$) as a function of the reward prediction update speed $\beta$.** Left: plotted on a linear scale. Right: plotted on a logarithmic scale. The theory (orange) matches the simulation data (blue), which show the average across 10,000 repetitions.

This can be considered to be the 'variance' caused by reward noise when there is 0 'bias' between the true expected reward and the reward prediction ($RP = \mathbb{E}[R]$). This variance function is illustrated in Figure S4 for $t = 1,000$. The squared RPE increases monotonically with $\beta$ and ranges between $\sigma^2$ and $2\sigma^2$. On a $Log_{10}$ scale, we can see that the squared RPE almost reaches its minimum value when the update speed $\beta$ is smaller than 0.1. The reason is straightforward. For any $\beta$ not close to 0, the term $(1-\beta)^{2(t-1)}$ is negligible when $t$ is large. Thus, the squared RPE is mainly affected by the term $\beta/(2-\beta)$, which takes its maximum value of 1 when $\beta = 1$ and is reduced to about 0.05 when $\beta = 0.1$ (and further reduced to about 0.005 when $\beta = 0.01$). As a result, the variance can be reduced approximately by half by choosing $\beta < 0.1$ (note that there is a constant $\sigma^2$ term that cannot be reduced).

**The bias-variance tradeoff**

We next consider a scenario in which the reward distribution is changing. During learning, the expected reward typically increases with time. For simplicity of calculations, we assume that the expected reward increases at a constant rate $\eta$:

$$\mathbb{E}[R_t] = (t-1)\eta, \tag{S86}$$

and the variance at all times equals $\sigma^2$.

When $\beta = 0$, the reward prediction $RP_t$ always remains 0. Then the expectation value of the squared reward prediction error is:

$$\mathbb{E}[(R_t - RP_t)^2] = Var[R_t - RP_t] + \mathbb{E}[R_t - RP_t]^2 \tag{S87}$$

$$= Var[R_t] + \mathbb{E}[R_t]^2 = \sigma^2 + (t-1)^2\eta^2. \tag{S88}$$

Note that the second term is the squared bias between the true expected reward and the reward prediction.

When $\beta = 1$, $RP_t = R_{t-1}$. In this case:

$$\mathbb{E}[(R_t - RP_t)^2] = \mathbb{E}[(R_t - R_{t-1})^2] = Var[R_t - R_{t-1}] + \mathbb{E}[R_t - R_{t-1}]^2 \tag{S89}$$

$$= 2\sigma^2 + \eta^2. \tag{S90}$$

Thus, using a fast update speed increases the variance term (from $\sigma^2$ to $2\sigma^2$) but reduces the bias term (from $(t-1)^2\eta^2$ to $\eta^2$).

In general:

$$\mathbb{E}[(R_t - RP_t)^2] = Var[R_t - RP_t] + \mathbb{E}[R_t - RP_t]^2 \tag{S91}$$

$$= Var[R_t] + Var[\sum_{i=0}^{t-2} \beta(1-\beta)^i R_{t-1-i}] \tag{S92}$$

$$+\Big(\mathbb{E}[R_t] - \mathbb{E}[\sum_{i=0}^{t-2} \beta(1-\beta)^i R_{t-1-i}]\Big)^2 \tag{S93}$$

$$= [1 + \frac{\beta}{2-\beta}[1-(1-\beta)^{2(t-1)}]]\sigma^2 \tag{S94}$$

$$+\Big(1 - \frac{(1-\beta)^{t-1} + (t-1)\beta - 1}{\beta(t-1)}\Big)^2 (t-1)^2\eta^2, \tag{S95}$$

where we have used the following formula in the derivation above:

$$\sum_{i=1}^{n} ia^i = \frac{na^{n+2} - (n+1)a^{n+1} + a}{(a-1)^2}. \tag{S96}$$

Comparing this result with the result obtained with a static reward (Eq. S85), we can see that the same 'variance' term remains, and there is a second term caused by the change of reward, which equals the square of the expected difference between the true expected reward and the reward prediction ($\mathbb{E}[R_t - RP_t]^2$) and thus can be considered as the 'bias'. The bias term decreases monotonically from $(t-1)^2\eta^2$ when $\beta = 0$ to $\eta^2$ when $\beta = 1$.

The squared RPE is illustrated in Figure S5. Overall, we can see that the squared RPE is a U-shaped function of $\beta$. If $\beta$ is close to 1, the squared RPE is large because of a large variance even though the bias is small. If $\beta$ is close to 0, the squared RPE is large due to a large bias even though the variance is small. A small squared RPE is achieved for an intermediate range of $\beta$ values (around $0.01 - 0.1$) for which the bias-variance tradeoff is well-balanced.

Is there an optimal update speed $\beta^*$ such that the squared RPE is strictly minimized? If $\sigma$ (in the variance term) dominates, the optimal update speed will be close to 0. If $(t-1)\eta$ (in the bias term) dominates, the optimal update speed will be close to 1. In general, one

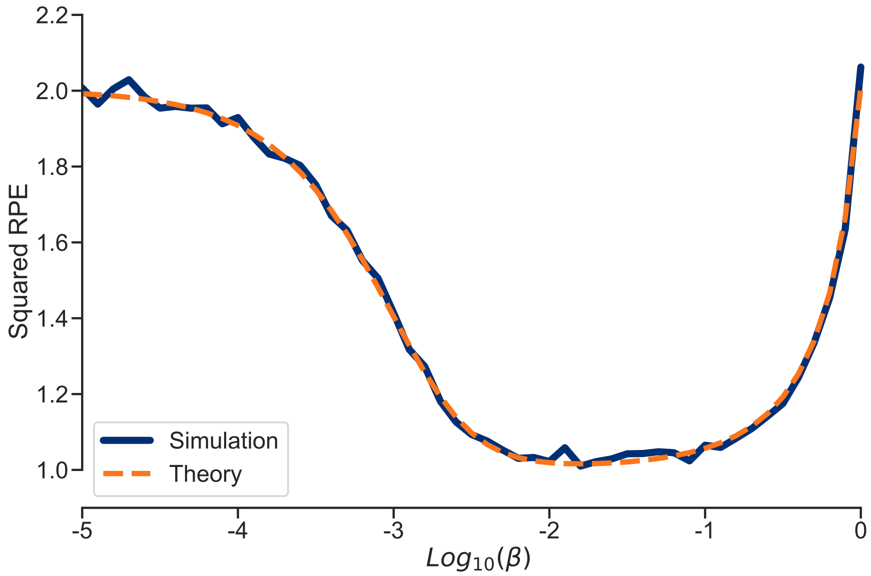


**Figure S5: Squared RPE as a function of $Log_{10}\beta$.** $\sigma = 1$, $\eta = 0.001$, $t = 1,000$. The theory (orange) matches the simulation data (blue), which show the average across $10,000$ repetitions.

can calculate the derivative of $\mathbb{E}[(R_t - RP_t)^2]$ with respect to $\beta$ and solve for the extreme point. However, $\beta^*$ will be a complicated function of $\sigma$, $\eta$, and $t$ and may not have a simple analytical form. Moreover, $\sigma$ and $\eta$ are generally unknown to the learning agent and adjusting the update speed as a function of $t$ may not be realistic. Thus, we take another approach by considering the coefficients of the variance and bias terms:

$$Squared\ RPE = Variance + Bias := A \times 2\sigma^2 + B \times (t-1)^2\eta^2, \tag{S97}$$

where $2\sigma^2$ is the maximum possible variance value (obtained when $\beta = 1$), $(t-1)^2\eta^2$ is the maximum bias value (obtained when $\beta = 0$), and the coefficients $A$ and $B$ are defined as:

$$A := \frac{1}{2}\Big[1 + \frac{\beta}{2-\beta}[1 - (1-\beta)^{2(t-1)}]\Big]. \tag{S98}$$

$$B := \Big(1 - \frac{(1-\beta)^{t-1} + (t-1)\beta - 1}{\beta(t-1)}\Big)^2. \tag{S99}$$

Figure S6 shows the two coefficients $A$ and $B$ as a function of $\beta$. We can see that the coefficient $A$ is sufficiently close to its minimum value of 0.5 when $\beta < 0.1$ (this is largely independent of trial number $t$). On the other hand, the coefficient $B$ is sufficiently close to its minimum value of $1/(t-1)^2$ when $\beta > 0.01$ for a sufficiently large number of trials ($t = 1,000$ shown).

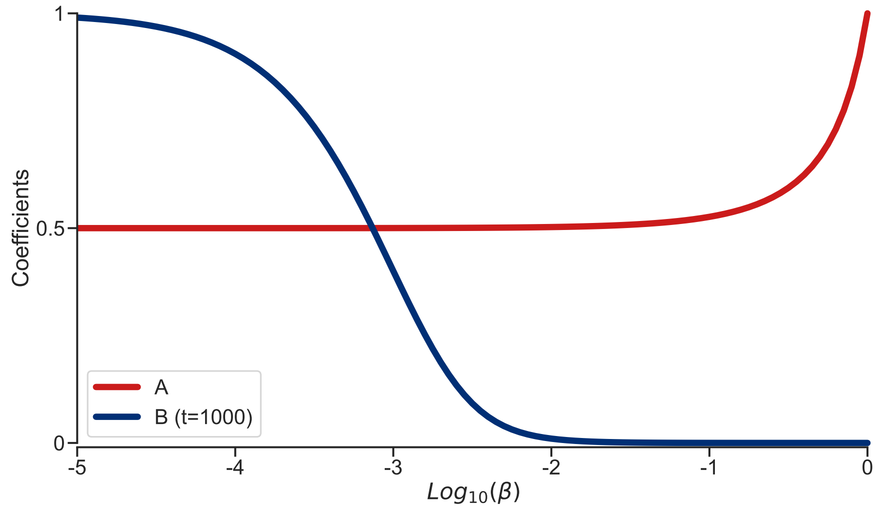


**Figure S6: Coefficients of variance and bias terms as a function of $Log_{10}\beta$.**

In conclusion, this analysis suggests that using $\beta < 0.1$ efficiently mitigates the variance and using $\beta > 0.01$ efficiently mitigates the bias, so that a range of $\beta = 0.01 - 0.1$ achieves a good bias-variance tradeoff. We note that these values that optimize the critic performance alone are much smaller than the update speed $\beta \approx 0.3$ found in natural song learning [4] and shown in the main text to maximize performance when the dynamics of learning are taken into account (Figure 7E).
